# HyperdCas12a-Based Multiplexed Genetic Regulation in *Candida albicans*

**DOI:** 10.1101/2025.10.20.683421

**Authors:** Nicholas C. Gervais, Ruby K.J. Rogers, Madeleine R. Robin, Rebecca S. Shapiro

## Abstract

Complex microbial phenotypes involve the combined activity of diverse gene regulatory networks. However, the majority of reverse genetics approaches in microbial pathogenesis research have focused on single-gene perturbation studies, in part due to the lack of available genetic tools in many pathogens. Developing enhanced versions of CRISPR-Cas platforms holds significant promise for improving the scalability of microbial functional genomics research. Here, we demonstrate highly efficient, inducible, and multiplexed activation and repression in the major human fungal pathogen *Candida albicans* by translating the hyperdCas12a variant to the fungal kingdom. This represents the first application of a CRISPR-Cas12 system in a human fungal pathogen. We profile the effectiveness of our new CRISPRa and CRISPRi tools and achieve tunable levels of target modulation. Further, we demonstrate that perturbing combinations of genes in the drug efflux and ergosterol biosynthesis pathways reveals important redundancies and synergistic properties in drug resistance circuitry. Our hyperdCas12a platform is thus an efficient system for the rapid generation of combinatorial mutants that will enable the mechanistic understanding of genetic interactions involved in diverse phenotypes in *C. albicans*. The enhanced activity with hyperdCas12a in fungi suggests it could be translated to other microbes as a powerful tool for studying genetic interactions.

## Introduction

Studying genetic interactions is critical for understanding how the functional relationships between genes collectively influence complex phenotypes [1]. Indeed, genetic interactions often underlie the phenotypic variability observed between individuals following a single mutation, as the genetic context in which a mutation arises will influence the resulting outcome [2]. The advent of new functional genomics technologies, whereby two or more genes can be mutated simultaneously, has facilitated an understanding of the critical and complex roles of genetic interactions in model systems [3]. For example, double gene deletion approaches have allowed for the thorough profiling of genetic interactions in the model yeast *Saccharomyces cerevisiae*, which have enabled the ordering of genetic pathways, the discovery of specific interactions important for growth in distinct environmental conditions, and the characterization of broader indirect roles that specific genes have on phenotypes [4,5]. These foundational studies have also helped categorize the disparate cellular consequences that genetic interactions have, whereby synthetic lethality [6], synergy [7], buffering [8], suppression [9], masking [10], and functional redundancy [11,12], for example, all represent different outcomes of an interaction. Additionally, genetic interactions are dependent on the expression levels of the corresponding genes, such that an interaction may switch from suppressive to enhancing following a change in the expression of one gene in the pair [13]. The spatial organization of the genome can also be dictated by genetic interactions, such as the non-random distribution of synthetic lethal pairs [14]. Genetic interactions may also influence the size of the genome, as functional redundancies of important cellular processes can be beneficial to the cell, thereby increasing the number of genes in the genome [15]. Thus, the biology of organisms is largely dictated by the sum effects of countless contributing genetic interactions.

To comprehensively investigate genetic interactions in diverse microbial species, new technological platforms must be developed [16]. Recent technological advances in model systems have enabled the identification of a myriad of genetic interactions and non-additive phenotypic outcomes caused by changes in two or more genes [17–21]. Applying such approaches in other microbial species will address several key limitations in current microbial research. For example, single mutations associated with drug resistance in clinical microbial isolates are often recapitulated *in vitro* and evaluated for their individual effects. However, many drug resistance phenotypes observed in clinical isolates and in laboratory-acquired strains are necessarily produced through epistatic relationships involving multiple genes [22,23]. Many of these genetic interactions related to drug resistance are also strain-dependent [24,25], and can drive the accelerated evolution of compensatory mutations compared to single-gene mutation-causing resistance [26]. Genetic tools that enable pairwise perturbation of genes may allow for the discovery of synthetic lethal combinations that could serve as drug targets for novel combination therapies [27,28], or targets that would re-sensitize multi-drug-resistant pathogens to existing antimicrobials [29]. Platforms that allow for the perturbation of essential genes are also highly desirable, as essential genes have significantly more interactions than non-essential genes [4]. This is of particular significance when studying microbial pathogens as essential genes are generally considered to be ideal targets for novel antimicrobials [30]. Harnessing platforms that allow for three or more genes to be altered is also critical, as trigenic interactions can be up to 100X more frequent than digenic interactions [31]. Thus, establishing new genetic tools that are capable of rapidly identifying genetic interactions in human pathogens will create many distinct opportunities for improving patient outcomes following infection.

While genetic interaction analysis has proven to be a powerful tool to identify epistatic interactions in diverse microbial species, this phenomenon is not well characterized in microbial fungi. This is of particular importance as fungal species have a significant global impact on human and plant health [32]. Human fungal pathogens are responsible for millions of deaths each year, and treatment failure is common in part due to the limited number of available antifungal therapeutics [33]. *Candida albicans* is the most common causative agent of human fungal infection, and hundreds of thousands of patients acquire invasive bloodstream *C. albicans* infections annually [34,35]. Increasing rates of drug resistance and drug tolerance in *C. albicans* have reduced the effectiveness of existing treatment options [36–38]. For example, while the azole class of antifungal drugs represent one of the most commonly employed therapeutic classes for invasive fungal infections, *C. albicans* can develop azole resistance via numerous distinct mechanisms [39]. While recent efforts to catalogue common mutations associated with antifungal drug resistance have helped highlight the influence of single-gene-encoded products [40], the phenotypic outcome of the convergence of multiple resistance mechanisms is more challenging to ascertain. New systems biology approaches will be necessary to begin uncovering the complex network of genetic interactions that contribute to diverse fungal phenotypes.

Generating double gene deletions via CRISPR-Cas9 has proven to be an effective strategy for dissecting genetic networks in drug efflux and virulence in *C. albicans* [41,42]. However, this approach is limited to studying two genes at once, and deletion-based systems inherently cannot be used to study essential genes or how interactions change due to varying levels of expression of the given gene(s) [13,43]. In addition, deletion-limited approaches will overlook genetic interactions caused by the overexpression of at least one of the involved genes [44,45]. Practically, constructing these mutants involves a relatively laborious protocol that requires synthesizing complex repair templates, making the process of building mutant libraries arduous and limiting the number of genes that can be included. Another CRISPR-based technique that has been extensively used to study genetic interactions involves the use of Cas12 [46–48]. CRISPR-Cas12 tools have been ubiquitously applied for multiplexed gene targeting, in part since, unlike Cas9, Cas12 proteins possess intrinsic RNase activity [49]. Thus, a single transcript containing a compact CRISPR array with CRISPR RNAs (crRNAs) that match the target regions that are interspersed by small direct-repeat (DR) regions can be processed by Cas12 into its individual constituents to target multiple distinct genomic sites [50,51]. CRISPR platforms with a nuclease-deficient version of the Cas protein (dCas) have additionally been extensively applied to repress (via CRISPR interference) or overexpress (via CRISPR activation) target gene expression [52]. Notably, as CRISPR-dCas tools allow for the reversible perturbation of essential genes, they often allow for more diverse genes to be studied than traditional active CRISPR-Cas systems [52]. Combining the advantages of Cas12 and dCas-based systems allows for multiplexed modulation of target gene expression [53]. Recent efforts to improve the efficiency of these platforms have further positioned dCas12-based tools to be powerful for studying complex genetic interactions [54,55].

Here, we develop the first CRISPR-dCas12a system in *C. albicans* for targeted and inducible overexpression or repression of multiple genes through simple multiplexed crRNA array construction. We demonstrate the translatability of the “hyper-efficient” d*Lb*Cas12a protein and its improved activation capacity in the fungal kingdom, allowing us to consistently drive higher levels of overexpression than previously, up to nearly 300-fold. We also expand the hyper*Lb*dCas12a system for targeted repression via CRISPR interference (CRISPRi), demonstrating levels of repression unattainable with our previous tools, and generate inducible versions of the CRISPR activation (CRISPRa) and CRISPRi systems. Finally, we use the new platform to identify functional redundancies in drug efflux upregulation, as well as massively synergistic resistance involving two genes in the ergosterol biosynthesis pathway. To our knowledge, our system is the first to allow for targeted overexpression or repression of multiple genes simultaneously in *C. albicans*. The simplicity with which these powerful new tools can be applied will provide new opportunities to study genetic interaction-based phenotypes at scale in *C. albicans*.

## Results

### CRISPRi/a Target Sites are More Accessible with dCas12a than dCas9 in C. albicans

To develop a novel CRISPR-dCas12 system for use in *C. albicans*, we first sought to assess the applicability of dCas12 for the *C. albicans* genome based on PAM site availability and how it differs from dCas9, for which CRISPR tools already exist in *C. albicans* [41,56,57]. When comparing the sum of all canonical Cas9 and Cas12a PAM sites across the entire genome, there appeared to be more PAM sequences targetable with Cas9 (**FIGURE 1A**), which aligns with our previous findings [58]. The PAM sites were approximately evenly distributed across each chromosome when normalizing by chromosome size (**SUPPLEMENTARY FIGURE 1**). To define the utility of dCas12 tools for modulating *C. albicans* gene expression, we additionally searched for PAM sites in the promoter region upstream of ORFs. Our previous attempts at comparing activation efficiency revealed that targeting upstream of the start codon was equal to or superior to targeting the TSS with currently available *C. albicans* TSS prediction data [57]. We therefore searched for PAM sites available only in a window 90bp-370bp upstream of start codons, and found a higher abundance for Cas12a than Cas9 (**FIGURE 1B**). Further, we found that, PAM sites were more evenly distributed between promoters for Cas12a than Cas9, as can be seen with a more extreme right skew for the distribution of Cas9 PAM (Cas9 D’Agostino Normality Test: statistic=512.053, p-value=6.44e-112; Cas12a D’Agostino Normality Test: statistic=70.991, p-value=3.84e-16) (**FIGURE 1C**). As a result, many of the Cas9 PAM sites in the genome are concentrated in a small number of promoters, whereas there are more promoters that have a modest number of PAM sites for Cas12a (**FIGURE 1D**). Generally, it is prudent to design several crRNAs for a given gene to ensure successful on-target activity occurs with at least one of them. Not only does Cas12a have more PAM sites available for modulating target gene expression overall, but it is also much more likely that designing a larger number of crRNAs will be possible with Cas12a for any given target gene, unless more than around 20 PAM sites are needed for a specific purpose, which would be unlikely (**FIGURE 1E**). Thus, dCas12a systems may better suited for effectively targeting promoter regions in the *C. albicans* genome. Regardless, combining the PAM sites of dCas9 with dCas12a will greatly increase the number of targetable sites in the genome overall.

**Figure 1.**
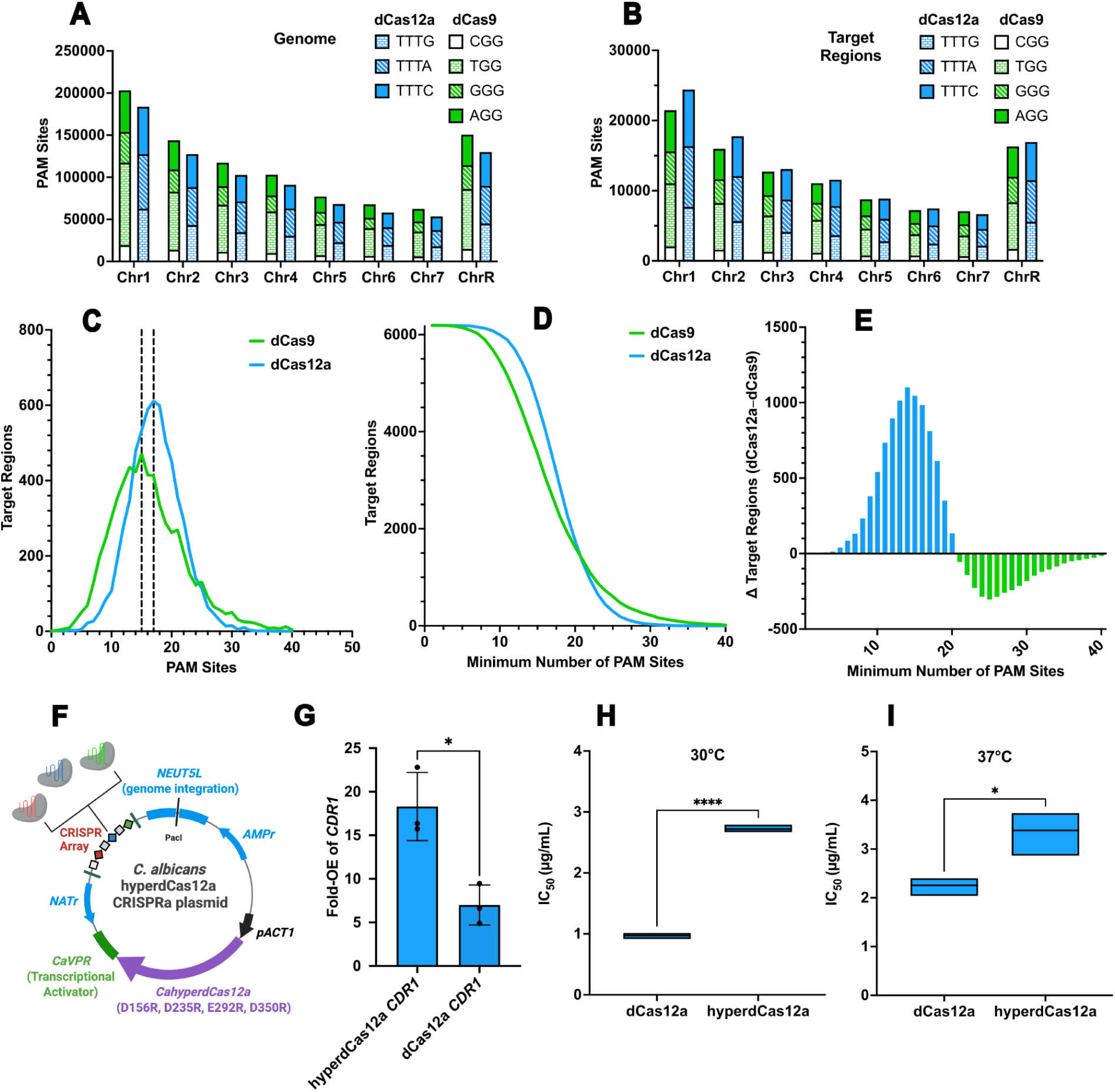
Implementation and advantages of CRISPR-dCas12a in *C. albicans*. (**A**) PAM sites were searched across the *C. albicans* genome by using a custom Python script. The total number for each Cas protein was identified and plotted per chromosome and delineated by the specific PAM recognition motif. (**B**) A region 90-370bp upstream of the start codon (‘Target Region’) for each of the ∼6000 canonical ORFs in the *C. albicans* genome was first isolated, and the corresponding PAM sites for both Cas proteins were counted and plotted similarly to in Figure 1A. (**C**) A distribution showing the number of Target Regions that have the corresponding number of PAM sites for both of the dCas9 and dCas12a proteins. The dotted line at the peak thus represents the most common number of PAM sites available for any given Target Region with each dCas protein. (**D**) A function that shows the number of Target Regions that have at least a certain number of PAM sites with dCas9 and dCas12a. (**E**) A differential function that shows how many more Target Regions have at least a certain number of dCas12a PAM sites than dCas9. For example, there are around 1000 more Target Regions that have at least 15 dCas12a PAM sites than there are Target Regions that have at least 15 dCas9 PAM sites. (**F**) A schematic of the hyperdCas12a CRISPRa system in *C. albicans*. The codon-optimized hyperdCas12a-VPR construct is expressed by the constitutively active *C. albicans ACT1* promoter. The *NEUT5L* sequence on the plasmid is homologous to the *NEUT5L* region on chromosome 5 in *C. albicans*, allowing for integration of the plasmid into the genome following plasmid linearization with the PacI enzyme. A CRISPR array with crRNAs interspersed by direct repeats can be cloned into the plasmid, and is expressed by the *C. albicans SNR52* promoter. Created in BioRender. (**G**) Comparison of the overexpression of *CDR1* achieved with the dCas12a and hyperdCas12a CRISPRa systems. Strains were grown at 37°C in YPD for ∼4h, followed by freezing and RNA extraction. RT-qPCR was performed on the samples, and fold-changes were calculated as a comparison between the listed CRISPRa strain and the NTC. Error bars represent the standard deviation of the corresponding replicates. Statistics were performed with an unpaired, two-tailed, parametric Welch’s t-test, \*\**p* < 0.01. (**H**) Differences in the corresponding fluconazole IC_50_ values at 30°C when dCas12a or hyperdCas12a is used to target *CDR1* for overexpression via CRISPRa. Strains were grown in an MIC assay at 30°C in YPD with a concentration gradient of fluconazole, and OD600 values were measured after 24 hours of growth. A logistic dose-response curve was then fitted to the normalized OD600 values from the MIC data to obtain the IC_50_ values. Statistics were performed with an unpaired, two-tailed, parametric Welch’s t-test, \**p* < 0.05, \*\*\*\**p* < 0.0001. (**I**) Differences in the corresponding fluconazole IC_50_ values at 37°C when dCas12a or hyperdCas12a is used to target *CDR1* for overexpression via CRISPRa. MIC assays and IC_50_ value calculations, along with statistics, were performed in the same way as in Figure 1H.

### The hyperdCas12a system allows for efficient targeted overexpression in C. albicans

To generate a single-plasmid CRISPRa-dCas12a system for *C. albicans*, we codon-optimized a nuclease-deficient dCas12a originally from *Lachnospiraceae bacterium* ND2006 that has been previously demonstrated for use in *Saccharomyces cerevisiae* [59], and combined it with the VPR activation complex we have previously used to activate gene expression in *C. albicans* and *Nakaseomyces glabratus* (**FIGURE 1F**) [57,60]. VPR is a tripartite activator complex that has demonstrated capacity for efficient activation in many organisms [61,62]. We placed the dCas12a-VPR construct under the control of the strong constitutive promoter from *ACT1*, similar to our previous *C. albicans* CRISPR-dCas systems [56,57]. To demonstrate the efficacy of the system in *C. albicans*, we cloned in a single crRNA targeting *CDR1*, which encodes a multidrug transporter often implicated in drug resistance [63]. When comparing this dCas12a-*CDR1* strain with a control strain containing a non-targetting CRISPR plasmid backbone, we found that the *CDR1* transcript increased by 7.0-fold (**FIGURE 1G**), suggesting the dCas12a-VPR CRISPRa system is able to robustly activate gene expression in *C. albicans*. We further monitored the minimum inhibitory concentration (MIC) of the dCas12a-*CDR1* strain to confirm that genotypic overexpression of *CDR1* would recapitulate phenotypic increases in antifungal drug resistance. Indeed, we found the corresponding IC_50_ in fluconazole was increased with both dCas12 and hyperdCas12a, and by a specific amount depending on the growth temperature **(SUPPLEMENTARY FIGURE 2)**. Nonetheless, we found that the IC_50_ was significantly higher for hyperdCas12a than dCas12a in all cases (**FIGURES 1H**, **1I**).

Next, to increase the levels of overexpression achievable with our system, we incorporated four amino acid mutations into the sequence of our dCas12a, based on mutations previously described in the “hyperdCas12a” variant, which has been employed in human cells [54]. Upon cloning the same *CDR1*-targeting crRNA into the *C. albicans* hyperdCas12a-containing plasmid, we found that overexpression of *CDR1* improved to 18.3-fold (**FIGURE 1G**), with a corresponding increase in the IC_50_ in fluconazole (**FIGURE 1H**, **1I**). This suggests that the increased efficacy observed with hyperdCas12a in human cell lines translates to fungi, and that our hyperdCas12a system can be exploited for highly effective gene activation in *C. albicans*. Given the high level of activation with hyperdCas12a, we decided to proceed with this system in all subsequent work.

### Multiplexed targeting allows for the overexpression of multiple genes and tunable overexpression

To determine the efficiency and on-target activity of our hyperdCas12a system, we tested several different crRNAs individually targeting two other efflux pump genes, *CDR2* and *MDR1*. We first designed and generated seven *C. albicans* strains with different crRNAs targeting *CDR2* (four crRNAs) or *MDR1* (three crRNAs). We found that all seven corresponding overexpression strains had increased resistance to the antifungal fluconazole, with a significant increase in the IC_50_ compared with the non-targetting CRISPRa control strain (**FIGURE 2A**, **FIGURE 2B**). A unique advantage of Cas12-based systems is the ability to easily achieve multiplexed targeting by using multiple crRNAs interspersed by short DR regions in the plasmid. We therefore further sought to determine if tiling the promoter of one gene with multiple different crRNAs could result in improved or synergistic activation of gene expression, as it has been shown in other species that well-spaced crRNAs targeting the same promoter can enhance activation with dCas9-CRISPRa and repression with dCas9-CRISPRi [64–67]. Therefore, we selected two crRNAs targeting *CDR2* (*CDR2* crRNA #2 and *CDR2* crRNA #4) that resulted in mild fluconazole-resistance phenotypes when used individually (**FIGURE 2A**), and combined them in one hyperdCas12a array (**FIGURE 2C**). While the two crRNAs individually led to a 12.2-fold and 66.2-fold overexpression of *CDR2*, respectively, combining them led to 275.9-fold overexpression (**FIGURE 2D**). Despite the marked increase in overexpression when using both crRNAs together, it was neither additive (12.2-fold x 66.2-fold = 807.6-fold), nor synergistic (>807.6-fold). Nevertheless, this demonstrates that the combination of multiple crRNAs can markedly increase gene activation in *C. albicans*. We also observed that the enhanced level of overexpression achieved while using both crRNAs together resulted in a significantly higher level of resistance to fluconazole in the corresponding strains (**SUPPLEMENTARY FIGURE 3**). This may be particularly useful in cases when the level of overexpression driven by one crRNA alone results in a very subtle phenotype that could otherwise be missed. Such crRNA multiplexing is a unique advantage of the Cas12a system, demonstrating the importance of this tool for studying phenotypes such as drug resistance in *C. albicans*.

**Figure 2.**
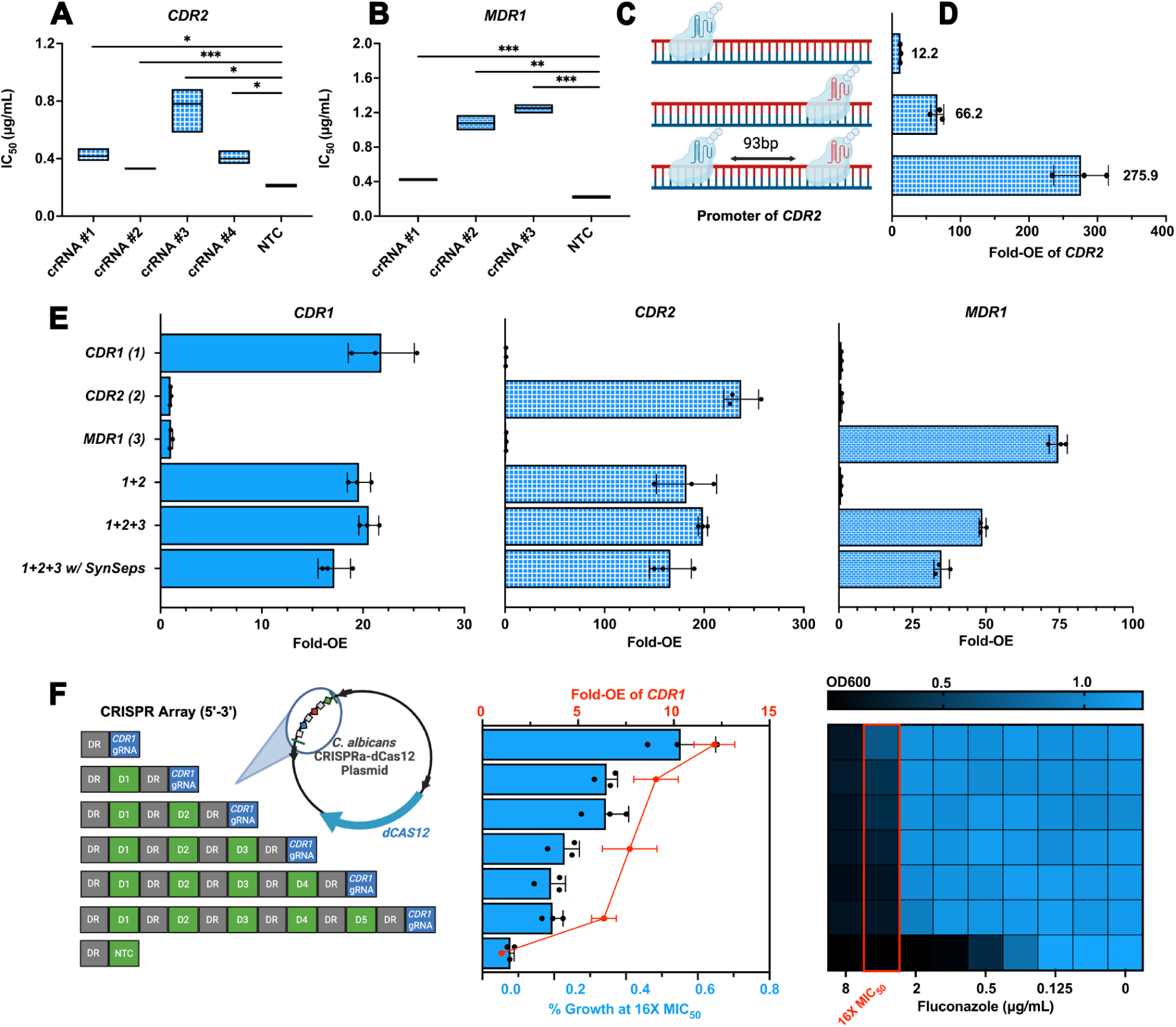
Validation of the multiplexed targeting capability of the hyperdCas12a system in *C. albicans*. (**A**) Comparison of the fluconazole IC_50_ values between different crRNAs used to target *CDR2* for overexpression via CRISPRa. Strains were grown in an MIC assay at 37°C in YPD with a concentration gradient of fluconazole, and OD600 values were measured after 24 hours of growth. A logistic dose-response curve was then fitted to the normalized OD600 values from the MIC data to obtain the IC_50_ values. Statistics were performed with an unpaired, two-tailed, parametric Welch’s t-test, \**p* < 0.05, \*\*\**p* < 0.001. (**B**) Comparison of the fluconazole IC_50_ values between different crRNAs used to target *CDR2* for overexpression via CRISPRa. The IC_50_ values were generated via MIC assays in the same way as for Figure 2A, and statistics were also calculated in the same way, \*\**p* < 0.01, \*\*\**p* < 0.001. (**C**) Schematic of how the hyperdCas12a system targets the *CDR2* promoter with only one of the two crRNAs as opposed to both at the same time. (**D**) Comparison of the overexpression of *CDR2* achieved in the singleplexed and multiplexed strains. Strains were grown at 37°C in YPD for ∼4h, followed by freezing and RNA extraction. RT-qPCR was performed on the samples, and fold-changes were calculated as a comparison between the listed CRISPRa strain and the NTC. Error bars represent the standard deviation of the corresponding replicates. (**E**) Comparison of the overexpression of *CDR1*, *CDR2*, and *MDR1* achieved in the singleplexed and multiplexed CRISPRa strains. The RT-qPCR experiment and statistics were performed in the same way as described for Figure 2D. (**F**) The impact of adding an increasing number of non-targeting crRNAs upstream of a *CDR1*-targeting crRNA on CRISPRa efficiency. The leftmost schematic (Created in BioRender) illustrates the structure of each CRISPR array corresponding to the expression and phenotypic data seen to the right, where green “DX” crRNAs correspond to different non-targeting crRNAs, and the “NTC” represents the array of the non-targeting control strain. To assess the corresponding level of overexpression of *CDR1*, strains were grown at 37°C in YPD for ∼4h, followed by freezing and RNA extraction. RT-qPCR was performed on the samples, and fold-changes were calculated as a comparison between the listed CRISPRa strain and the NTC strain. Error bars represent the standard deviation of the corresponding replicates. To determine the drug susceptibility profiles of each strain, fluconazole MIC assays were performed, where strains were grown at 37°C in YPD with a concentration gradient of fluconazole, and OD600 values were measured after 24 hours of growth. The OD600 values from the wells corresponding to a concentration of fluconazole 16X that of the MIC_50_ of the NTC strain were then overlaid onto the RT-qPCR data. Here, errror bars represent the standard deviation of the corresponding replicates. The % growth at the 16X MIC_50_ fluconazole concentration for each strain was calculated by normalizing the values of each strain at 16X the MIC_50_ of fluconazole for the NTC (ie. 4μg/mL) to the corresponding no-drug control well.

Next, to demonstrate whether the system could be used for multiplexed overexpression of multiple different genes simultaneously, we combined a *CDR1* crRNA in a single CRISPR array along with *CDR2* crRNA #3 and *MDR1* crRNA #3, as they led to the highest respective increase in the IC_50_, and thus presumably the highest level of overexpression. Indeed, targeting all three of the efflux pump genes together led to the overexpression of all three (**FIGURE 2E**). Importantly, the basal expression of each gene did not seem to be influenced by the increased expression of one of the other genes in the individual overexpression strains (**FIGURE 2E**). Therefore, the achieved overexpression in the multiplexed strains is due to the activity of the corresponding crRNA, and not a result of changes in how the gene was regulated in response to the overexpression of one of the other genes. We did find, however, that the level of overexpression achieved for some genes in the multiplexed strains seemed to be slightly lower than when the corresponding gene was targeted individually, particularly for *MDR1* (**FIGURE 2E**), whose representative crRNA was in the 3’-most position in the CRISPR array. The activity of downstream crRNAs can sometimes be negatively affected by the presence of upstream crRNAs with specific sequences [55]. While high GC upstream crRNAs are most likely to interfere with the activity of downstream crRNAs [55], and neither the *CDR1* crRNA nor *CDR2* crRNA have a particularly high GC content, we nonetheless decided to test whether the addition of synthetic separators (*synSeparators*) upstream of each direct repeat (DR) would help restore the level of overexpression of *MDR1*. Interestingly, the addition of the *synSeparators* appeared to further lower the level of overexpression of *MDR1* (**FIGURE 2E**), thus suggesting that the reduced overexpression of *MDR1* in the multiplexed strain is not due to direct interference from the upstream *CDR1* and *CDR2* crRNAs. The slightly lower activation efficiency achieved for all three target genes with the *synSeparators* implies that they were not beneficial in this particular instance.

It has previously been shown with diverse CRISPR systems that while the position of the crRNA in the array does not seem to affect its activity [55,68], an increased number of crRNAs in the array reduce the activities of each individual crRNA [68]. We therefore decided to characterize how the level of overexpression achieved would be affected upon adding an increasing number of non-targeting crRNAs upstream of a working *CDR1*-targeting crRNA. Even when there were five non-targeting crRNAs upstream of the *CDR1* crRNA, we found that *CDR1* was still overexpressed by 6.3-fold (**FIGURE 2F**). As the strain with the single *CDR1-*targetting crRNA had a fold-OE of *CDR1* of 12.1, the level of overexpression of *CDR1* was therefore reduced by around 1.9-fold with the presence of the 5 non-targeting crRNAs upstream in the array (**FIGURE 2F**). The buffered level of overexpression of *CDR1* by adding non-targeting crRNAs into the array suggests that this could serve as a useful alternative strategy for tuning the level of overexpression produced with the system. Interestingly, we found that while the reduced overexpression of *CDR1* correlated with a decrease in growth at 16X the MIC_50_ of fluconazole, the overall level of drug resistance in the corresponding strains was fairly consistent across all strains, and thus scaled non-linearly with the overexpression of *CDR1* (**FIGURE 2F**). This suggests that reaching a certain threshold of overexpression of *CDR1* results in a large increase in fluconazole resistance, but with diminishing returns in further levels of drug resistance following additional upregulation of *CDR1*. The successful overexpression of *CDR1* along with the addition of non-targeting crRNAs in the array suggests that our system can be used for multiplexed CRISPR array processing, and that the number of crRNAs in the array may modestly affect the on-target activity of the system for the corresponding gene.

### The hyperdCas12a platform is efficient for multiplexed gene repression and inducible gene regulation

Given the ability for hyperdCas12a CRISPRa to robustly overexpress genes, singly and in combination, we next aimed to assess whether hyperdCas12a could similarly be used for CRISPRi-based genetic repression. Repressing the expression of many genes in the same cell can enable genetic interaction analysis and help uncover and order the genetic pathways underpinning diverse cellular processes. Targeted repression can also complement genetic overexpression, as increasing the expression of a gene often does not simply result in an opposite phenotype as may be expected by decreasing its expression [50,69]. Therefore, we next replaced the transcriptional activator complex VPR with two copies of the *C. albicans* codon-optimized version of the mammalian repressor Mxi1, as we have previously exploited in our CRISPRi-dCas9 system [56]. To demonstrate the efficacy of our hyperdCas12a-CRISPRi system, we targeted *ERG6* and *ERG251*, two members of the ergosterol biosynthesis pathway, individually and in combination. We found that both genes were highly repressed with this system, with *ERG6* up to 18.4-fold (**FIGURE 3A**), and *ERG251* up to 28.0-fold (**FIGURE 3B**), demonstrating the capacity of hyperdCas12a-CRISPRi to effectively and strongly repress genes in *C. albicans*. Next, we tested multiplexed repression when targeting *ERG6* and *ERG251* simultaneously, and found we were able to effectively repress both genes, though here the levels of repression were slightly lower at 6.0-fold (**FIGURE 3A**) and 11.8-fold (**FIGURE 3B**), respectively, suggesting that the system was more sensitive to the presence of multiple crRNAs in this situation.

**Figure 3.**
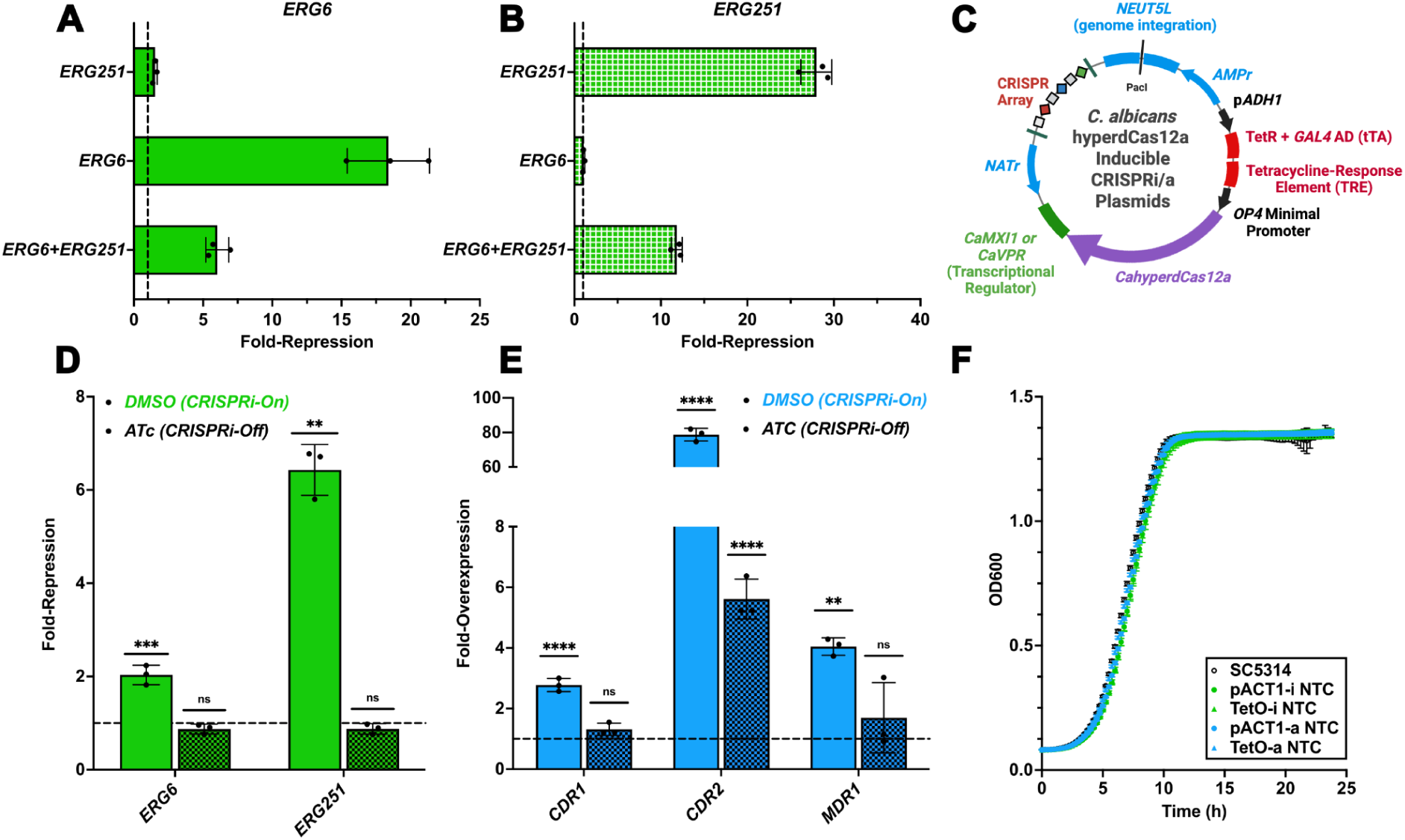
Expansion of the hyperdCas12a toolkit in *C. albicans* to include a highly efficient CRISPRi system as well as inducible versions of both the CRISPRa and CRISPRi platforms. (**A**) Comparison of the repression of *ERG6* achieved in the singleplexed and multiplexed CRISPRi strains. Strains were grown at 37°C in YPD for ∼4h, followed by freezing and RNA extraction. RT-qPCR was performed on the samples, and fold-changes were calculated as a comparison between the listed CRISPRi strain and the corresponding NTC strain. Error bars represent the standard deviation of the corresponding replicates. (**B**) Comparison of the repression of *ERG251* achieved in the singleplexed and multiplexed strains. *ERG251* gene expression changes for all strains were monitored in the same samples and at the same time as in panel 4A. (**C**) Schematic of the inducible promoter system added to both the CRISPRi and CRISPRa hyperdCas12a systems in *C. albicans*. The *ADH1* promoter from *C. albicans* drives expression of the tetracycline repressor protein (TetR) fused to a *GAL4* activation domain (together known as the tetracycline-controlled transactivator [tTA]), which binds to the tetracycline-response element (TRE) in the absence of tetracycline, thereby activating expression of hyperdCas12a from the *OP4* minimal promoter. In the presence of tetracycline, the tTA is released from the TRE, thereby inactivating expression of hyperdCas12a. Created in BioRender. (**D**) Comparison of the repression of *ERG6* and *ERG251* achieved in the multiplexed CRISPRi strain in the presence of vehicle (DMSO) alone and tetracycline. Strains were grown at 30°C in YPD with 250ng/mL ATc overnight, then diluted in fresh YPD with 250ng/mL ATc and grown for ∼5h at 37°C, followed by freezing and RNA extraction. RT-qPCR was performed on the samples, and fold-changes were calculated as a comparison between the listed CRISPRi strain and the corresponding NTC strain. Error bars represent the standard deviation of the corresponding replicates. Error bars represent the standard deviation of the corresponding replicates. Statistical differences between when the system was on (DMSO) and off (ATc) compared to the same growth conditions for the NTC are shown, and were calculated with an unpaired, two-tailed, parametric Welch’s t-test, \*\*\**p* < 0.001, \*\*\*\**p* < 0.0001. (**E**) Comparison of the overexpression of *CDR1*, *CDR2*, and *MDR1* achieved in the multiplexed CRISPRa strain in the presence of vehicle (DMSO) alone and tetracycline. RNA extractions and RT-qPCR experiments were performed in the same way as in Figure 3D, ns = *p* > 0.05, \*\**p* < 0.01, \*\*\*\**p* < 0.0001. (**F**) Kinetic growth curve assay in nutrient-rich media with the constitutive and inducible NTC hyperdCas12a strains. Strains were grown at 37°C in YPD for 24h, where the OD600 value was measured every 15 minutes and plotted as a function of time. Error bars represent the standard deviation of the corresponding replicates.

Next, we sought to determine if we could generate a regulatable version of the hyperdCas12a systems to enable controlled regulation of gene overexpression and repression in *C. albicans.* Temporal control of CRISPR-mediated genomic alterations can be desirable when studying the function of a gene at a specific growth period, when the target genes are essential, or when constitutive activity of the system would result in cell death, for example [70]. Therefore, we next replaced the *ACT1* promoter driving the expression of the dCas12a fusion protein in both our CRISPRa and CRISPRi plasmids with a tetracycline-inducible (Tet-Off) promoter system previously optimized for use in *C. albicans* [71]. We opted for a Tet-Off system, as opposed to a Tet-On system [72], as it does not require the inducing drug (tetracycline) to be present when the system is on, and we can therefore avoid situations where the tetracycline may interfere with our screening conditions, for instance by affecting the activity of antifungal drugs [73,74]. Our system utilizes a tetracycline repressor protein (TetR) fused to a *GAL4* activation domain, together known as the tetracycline-controlled transactivator (tTA) and driven by the *C. albicans ADH1* promoter, that binds to the Tetracycline Response Element (TRE) and drives expression of the dCas12a fusion protein from the *C. albicans OP4* minimal promoter in the absence of tetracycline (**FIGURE 3C**). When tetracycline is added, the tTA unbinds from the TRE, resulting in abrogated expression of dCas12a. To demonstrate its functionality for turning off the targeted multiplexed regulation, we cloned the *ERG6+ERG251* array used previously into the inducible CRISPRi plasmid, and the *CDR1+CDR2+MDR1* array used previously into the inducible CRISPRa plasmid. The addition of anhydrotetracycline hydrochloride (ATc) during growth completely shut off the significant repression of *ERG6* and *ERG251* that was caused by the CRISPRi system in the presence of the vehicle (DMSO) alone (**FIGURE 3D**). Similarly, ATc effectively shut off the expression of *CDR1*, *CDR2*, and *MDR1* in the multiplexed CRISPRa strain (**FIGURE 3E**), suggesting the regulatable Cas12-CRISPRi and CRISPRa systems can effectively alter gene expression in an inducible manner. While the level of *CDR2* overexpression was reduced 14-fold in the presence of ATc compared to vehicle alone, there was still 5.6-fold overexpression of *CDR2* compared to the non-targeting control (**FIGURE 3E**). As this crRNA resulted in the strongest activity of the system we have seen thus far, it is possible that extremely efficient crRNAs will result in some activity of the system even in the presence of ATc. This suggests that additional time growing in the presence of ATc may be required to fully prevent any gene expression changes caused by the system. It also demonstrates how the inducible systems may be used to tune the level of target gene expression modulation. Together, this demonstrates that our regulatable dCas12 CRISPRi and CRISPRa systems can be effectively used to temper the level of multiplexed overexpression or repression of target genes.

We further wanted to confirm that constitutive or regulatable expression of dCas12a did not significantly impact fungal fitness. Thus, we performed a kinetic growth-curve assay with four non-targetting dCas12a strains (representing both the CRISPRi and CRISPRa constitutive and inducible systems), along with the wild-type SC5314 strain, in plain nutrient-rich media for 24h at 37°C. We did not observe any differences in growth rate, carrying capacity, or lag time, suggesting that the simple integration of the empty vectors into the genome and the expression of the hyperdCas12a protein and a non-targeting crRNA does not result in any obvious growth defect (**FIGURE 3F**).

### Multiplexed targeting reveals diverse genetic interactions in C. albicans drug resistance pathways

Finally, given the capacity for dCas12a to efficiently repress or overexpress combinations of *C. albicans* genes, we sought to verify whether our system could be used to investigate the phenotypic contribution of different genes with overlapping roles, and reveal potential redundancies or synergies in genetic interaction networks. We examined genetic interactions mediating antifungal resistance phenotypes, focusing on overexpressing *CDR1*, *CDR2*, and *MDR1* with CRISPRa, as they all encode multidrug transporters associated with resistance to azoles. We also repressed *ERG6* and *ERG251* with CRISPRi due to their involvement in the biosynthesis of ergosterol, the target of azoles. To determine the phenotypic consequences of modulating these combination of genes, we assessed the levels of resistance and tolerance of our CRISPRa efflux pump strains to fluconazole (**FIGURE 4A**), and found that neither the IC_50_ (‘resistance’, **FIGURE 4B**) nor supra-MIC_80_ growth (‘tolerance’, **FIGURE 4C**) achieved via the overexpression of *CDR1* in combination with either *CDR2* or both *CDR2* and *MDR1* was higher than that of *CDR1* overexpression alone. This suggests that there is functional redundancy in the fluconazole drug efflux network in *C. albicans*, and that there may be a limit to how much the cell can decrease susceptibility to fluconazole through enhanced drug efflux alone.

**Figure 4.**
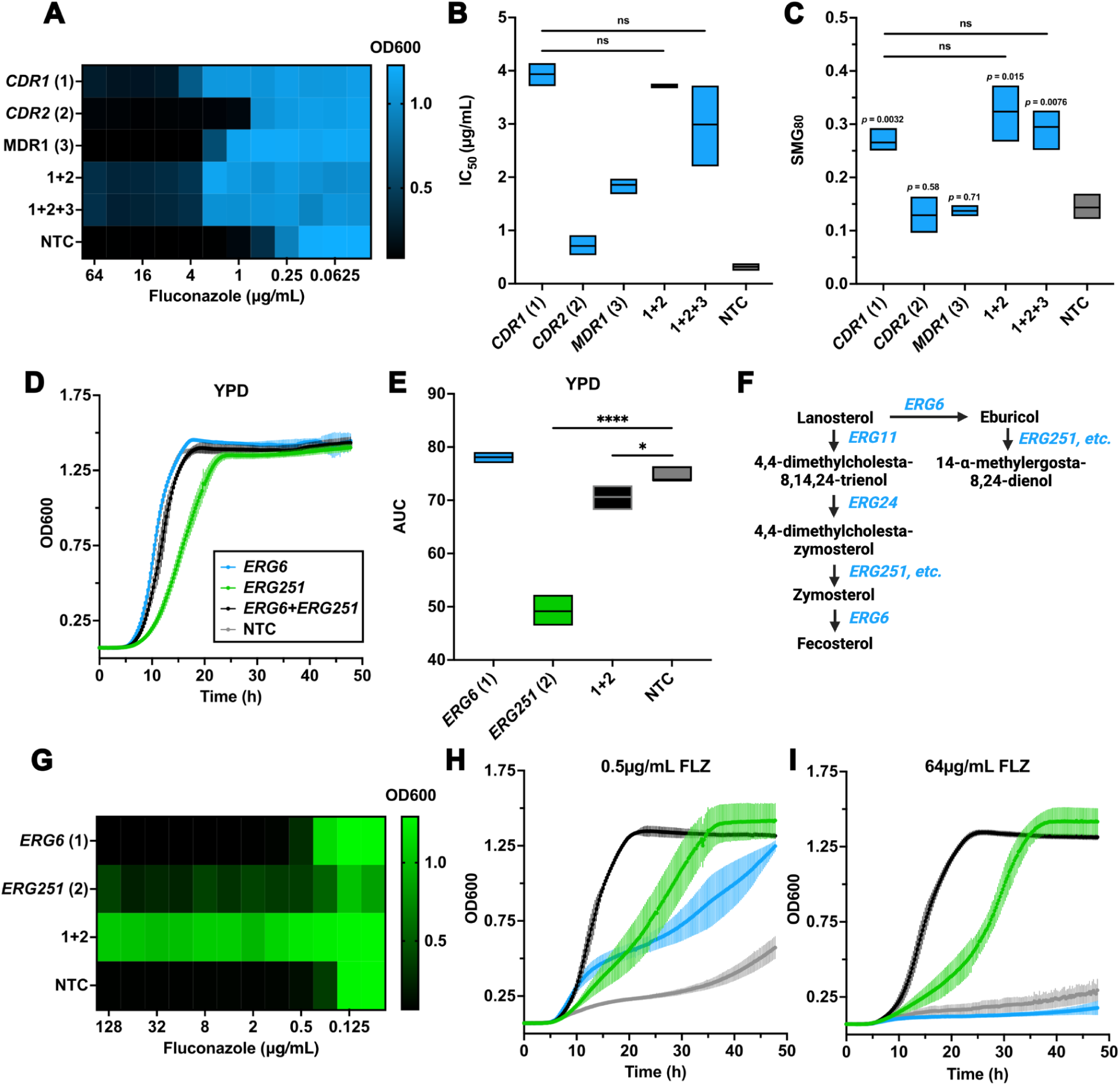
Characterization of diverse genetic interactions in the *C. albicans* drug efflux network and ergosterol biosynthesis pathway via hyperdCas12a. (**A**) Fluconazole MIC assay with the singleplexed and multiplexed drug efflux CRISPRa strains. Strains were grown at 37°C in YPD with a concentration gradient of fluconazole, and OD600 values were measured after 24 hours of growth. (**B**) Fluconazole IC_50_ values of the drug efflux CRISPRa strains. A logistic dose-response curve was fitted to the normalized OD600 values from the MIC assay in Figure 1A to obtain the IC_50_ values. Statistics were performed with an unpaired, two-tailed, parametric Welch’s t-test, ns *= p* > 0.05. (**C**) Fluconazole SMG values of the drug efflux CRISPRa strains. SMG values were calculated from the average growth of the given strain in the corresponding concentrations of drug above the MIC_80_ based on the OD600 values from the MIC assay in Figure 1A. Statistics were performed with an unpaired, two-tailed, parametric Welch’s t-test, ns *= p* > 0.05. (**D**) Area under the curve (AUC) values of the kinetic growth-curve assay shown in Figure 4. AUC values were calculated by using the trapezoidal rule with a custom Python script. Statistics were performed with an unpaired, two-tailed, parametric Welch’s t-test, ns *= p* > 0.05, \**p* < 0.05, \*\*\*\**p* < 0.0001. (**E**) Kinetic growth curve assay in nutrient-rich media with the singleplexed and multiplexed ergosterol biosynthesis CRISPRi strains. Strains were grown at 37°C in YPD for 24h, where the OD600 value was measured every 15 minutes and plotted as a function of time. Error bars represent the standard deviation of the corresponding replicates. (**F**) A simplified version of part of the main and alternate ergosterol biosynthesis pathway in *C. albicans*. Created in BioRender. (**G**) Fluconazole MIC assay with the singleplexed and multiplexed ergosterol biosynthesis CRISPRi strains. Strains were grown at 37°C in YPD with a concentration gradient of fluconazole, and OD600 values were measured after 24 hours of growth. (**H**) Kinetic growth curve assay in 0.5μg/mL of fluconazole with the singleplexed and multiplexed ergosterol biosynthesis CRISPRi strains. Strains were grown at 37°C in YPD supplemented with 0.5μg/mL of fluconazole for 48h, where the OD600 value was measured every 15 minutes and plotted as a function of time. Error bars represent the standard deviation of the corresponding replicates. The key in this panel is shared with Figure 4D, where the blue line represents the *ERG6* CRISPRi strain, the green line represents the *ERG251* CRISPRi strain, the black line represents the *ERG6*+*ERG251* CRISPRi strain, and the grey line represents the NTC strain. (**I**) Kinetic growth curve assay in 64μg/mL of fluconazole with the singleplexed and multiplexed ergosterol biosynthesis CRISPRi strains. The assay and statistics were performed in the same way as described for Figure 4H.

To further determine the phenotypic consequences of repressing combinations of genes, we measured drug resistance in *ERG* CRISPRi strains. We found that individually repressing *ERG251* led to a growth defect (**FIGURES 4D**, **4E**), which has been previously reported when *ERG251* is deleted [75]. Interestingly, the additional repression of *ERG6* in our multiplexed strain suppressed the growth defect and almost completely restored normal growth in nutrient-rich media (**FIGURES 4D**, **4E**). *ERG6* and *ERG251* are either directly upstream or downstream of each other in the primary ergosterol biosynthesis pathway and alternate pathway (**FIGURE 4F**) [75]. Therefore, repressing one may influence phenotypes caused by the repression of the other. Strikingly, we also observed that while the repression of *ERG6* alone led to only a modest increase in fluconazole resistance (**FIGURES 4G**, **4H**, **4I**), and the repression of *ERG251* alone led to an increase in fluconazole tolerance (**FIGURES 4G**, **4H**, **4I**), the combined repression of both led to a massive increase in the level of resistance (**FIGURES 4G**, **4H**, **4I**). Together, this work demonstrates that our novel hyperdCas12a CRISPR-dCas platform can be used to study the involvement of multidrug efflux and ergosterol biosynthesis in drug resistance. This represents a powerful tool for interrogating complex genetic circuitry that could be used to investigate countless other pathways underlying fungal phenotypes in the future.

## Discussion

In this work, we developed the first CRISPR-Cas12 platform for use in a major human fungal pathogen, and validated its ability to efficiently and combinatorially modulate target gene expression. We demonstrated that hyperdCas12a-CRISPRa can be used for a high level of gene overexpression, and tiling a gene promoter with multiple crRNAs can boost the level of achieved overexpression, resulting in an improved signal of phenotypes that may otherwise be overlooked (**SUPPLEMENTARY FIGURE 3**). We further show that inserting non-targeting crRNAs upstream of the targeting crRNA in the CRISPR array represents an effective strategy for tuning the level of on-target activity. We also expanded the system to allow for targeted gene repression via CRISPRi, and demonstrated that both activation and repression can be reversibly downregulated or fully silenced using the Tet-Off system. HyperdCas12a thus represents one of the only tools for targeted multiplexed overexpression or repression in *C. albicans*, and offers major improvements to existing CRISPR-dCas tools for single-gene perturbation in *C. albicans*. Using this optimized system, we constructed and profiled the first targeted multiplexed overexpression and repression mutants in *C. albicans*, which revealed striking genetic interactions. The combinatorial CRISPRa mutants revealed functional redundancies in the drug efflux network, indicating that drug efflux alone may not be sufficient to achieve high levels of resistance. Conversely, the multiplexed CRISPRi mutant revealed strong synergistic drug resistance and genetic suppression when the corresponding two members of the ergosterol biosynthesis pathway are repressed simultaneously. Overall, we have developped and optimized a highly-efficient and comprehensive CRISPR-dCas12 toolbox that will allow for further scalable investigation into the complex gene networks that underlie diverse fungal phenotypes.

The ability to efficiently and inducibly overexpress or repress combinations of genes using our hyperdCas12a system addresses many current limitations in functional genomics approaches for *C. albicans*, in particular for dissecting genetic interactions. Large-scale deletion libraries have been instrumental in defining the roles of many genes under diverse environmental stress conditions [76,77]. However, integrating these insights with an understanding of the mechanistic relationships between genes will require more flexible techniques. Previous efforts to characterize genetic interactions in *C. albicans* via chemical-genetic profiling or double-deletion approaches, while valuable, have been limited in scope and scalability [41,78]. In model systems, recent advances in CRISPR-Cas12 technologies have enabled the construction of large combinatorial perturbation libraries to study genetic interactions [47,48]. By establishing this hyperdCas12a system in *C. albicans*, we thus provide a platform to translate these powerful approaches to fungal pathogens. This system therefore holds promise for the systematic, high-throughput genetic interaction mapping in *C. albicans*, offering unprecedented opportunities to interrogate gene networks that underpin fungal adaptation, pathogenesis, and drug resistance. One scenario where such tools can be uniquely applied to study complex multi-gene interactions in *C. albicans* is in the study of aneuploidy-associated phenotypes. Aneuploidies, which frequently arise in *C. albicans* and are associated with antifungal drug resistance, often result in the differential regulation of hundreds of genes, with phenotypes commonly arising from the combined upregulation of multiple genes on trisomic chromosomes [79–83]. The novel hyperdCas12a tools presented here offer a novel approach for rapidly characterizing such genetic interactions in fungal genomes at scale.

As a tool for perturbing gene function in *C. albicans*, hyperdCas12a enables more efficient and accessible targeting. While only canonical PAM sites for both dCas9 and dCas12a were included in our analysis, hyper*Lb*dCas12a outperforms WT *Lb*dCas12a even with non-canonical PAM sites (eg. TTTT, CTTA, TTCA, TTCC) [54], implying that the potential availability of PAM sites for our *C. albicans* hyperdCas12a system may be much greater than shown here. Further, our observation that the improved activation capacity observed with hyperdCas12a translates to the fungal kingdom suggests a possible universal improvement for CRISPR-dCas12a systems across the tree of life. This may be particularly useful when attempting to develop CRISPR-dCas tools in intractable organisms where the genotypic and phenotypic readouts attainable with WT dCas proteins are weak, and have either precluded the meaningful discovery of important phenotypes, or have required an impractical number of crRNAs to be designed to accomplish successful genetic regulation. This is further highlighted by our ability to enhance activation by tiling the promoter with more than one crRNA, suggesting that only one CRISPR array needs to be designed and tested to determine how well the target gene could be modulated. Here, we showed that using two crRNAs targeting different regions of the *CDR2* promoter improved the level of activation, though not to an additive or synergistic extent. However, a gene with a lower level of basal expression may be subject to relatively higher levels of overexpression with CRISPRa [61], thus implying that additive or synergistic activation between two crRNAs may be possible in other situations.

It has long been established that the deletion of combinations of *CDR1*, *CDR2*, and *MDR1* progressively leads to increased fluconazole susceptibility [84]. Our current work, however, demonstrates that the overexpression of *CDR1* alone phenocopies the overexpression of *CDR1*, *CDR2*, and *MDR1* together (**FIGURE 4A**). This suggests that gene deletion may in some cases be a poor proxy for understanding the role of different genes in fungal phenotypes, and of the natural gain-of-function mutations that occur during the development of drug resistance in *C. albicans*. The subtle non-significant decreases observed between the strains may be due to an established genetic interaction between *CDR1* and *CDR2*, resulting in the fitness and drug resistance of the corresponding strain being lower than expected when the two genes are deleted together [41]. Consistent with previous findings, we observed with our CRISPRi system that the repression of *ERG251* led to a growth defect [75]. It has been suggested that this may in part be due to disrupted ergosterol production and the resulting accumulation of 4-methyl episterol (“sterol A”) and negative feedback on farnesol production [75]. We additionally found that repressing *ERG251* and *ERG6* together rescued the growth defect. The sterol eburicol would likely accumulate with the repression of *ERG251* in *C. albicans*, but would instead be returned to normal levels with the additional repression of the direct upstream producer of eburicol in the alternative pathway, *ERG6* (**FIGURE 4F**) [75]. It has previously been shown in the pathogenic mold *Aspergillus fumigatus* that accumulation of eburicol may be toxic to the fungal cell [85]. Therefore, the growth defect resulting from *ERG251* depletion may be at least in part due to the accumulation of eburicol. Inconsistent with previous findings, we did not observe any change in the expression of *ERG6* in our *ERG251* repression strain when grown in nutrient-rich media (**FIGURE 3A**) [75]. We also observed a strong synergistic effect that repressing both *ERG251* and *ERG6* had on resistance to fluconazole (**FIGURE 4G**). Previous work has shown that *ERG6* and *ERG251* are the two most downregulated genes in the ergosterol biosynthesis pathway upon fluconazole exposure when the *TLO* gene family, involved in global transcriptional control, is deleted in *C. albicans* [86]. Notably, the deletion of the *TLO* family also leads to reduced susceptibility to fluconazole [86]. Therefore, it is possible that our combined repression of *ERG6* and *ERG251* partly phenocopies the *TLO* mutant, leading to the accumulation of several sterol intermediates, including lanosterol, which has been suggested to contribute to its decreased drug susceptibility (**FIGURE 4F**) [86].

Since its original development in human cells, the modifications in hyperdCas12a have been demonstrated to have far-reaching benefits in the context of other synthetic biology tools. Indeed, new research suggests that hyperdCas12a is an effective platform for epigenome editing and for the characterization of large coding and non-coding regions of DNA via high-throughput combinatorial CRISPR screens [87]. Tools such as hyperdCas12a with the precision required to investigate non-coding DNA and other non-canonically functional elements of the genome could open up countless opportunities for understanding complex non-protein-driven fungal phenotypes at scale [88]. While our new hyperdCas12a toolbox promises the ability to comprehensively profile genetic interactions involving overexpression and repression in fungal genomes, these dCas12 systems could also be converted back to active Cas12, allowing for multiple deletions to be made in the same cell. An active Cas12 system of this kind capable of introducing multiple targeted double-stranded breaks in the cell could potentially be harnessed to simultaneously generate multiple deletions or loss-of-heterozygosity (LOH) events, for example, and study their compounding contributions to fungal fitness [58]. Additionally, as our hyperdCas12a platform represents one of the first non-*Sp*dCas9-based CRISPR tools to be employed for use in a major human fungal pathogen, it may now therefore be possible to harness both dCas9 and dCas12a together to perform simultaneous activation and repression in the same cell, similar to using orthologs of dCas12 proteins with different crRNA preferences [87]. Our hyperdCas12a toolbox, therefore, represents a highly efficient platform that can now be harnessed for dissecting genetic interactions temporally and at scale in *C. albicans*.

## Materials and Methods

### PAM Site Comparison

To assess the number of corresponding PAM sites available for dCas9 and dCas12a, the reference *C. albicans* SC5314 Assembly 22 genome was first downloaded from the *Candida* Genome Database [89]. PAM sites from the “A” haplotype alone were counted since we designed our crRNAs against haplotype A promoters. The PAM site sequences searched for dCas9 were CGG, TGG, GGG, and AGG (along with their respective reverse complements), and for dCas12a were TTTG, TTTA, and TTTC (along with their respective reverse complements).

### Media and growth conditions

*Candida albicans* (SC5314 background) strains were grown routinely at either 30°C or 37°C in yeast peptone dextrose (YPD) broth or plates supplemented with 250μg/mL nourseothricin (NAT) from Jena Biosciences (cat. AB-102L) for plasmid selection. 5-alpha Competent *Escherichia coli* cells from NEB (cat. C2987H) were grown at 30°C in Lysogeny Broth (LB) or plates supplemented with both 100μg/mL ampicillin (AMP) and 250μg/mL NAT for plasmid selection.

### Plasmid design and construction

All gene fragments and primers used for cloning are listed in **SUPPLEMENTARY TABLE 1** and **SUPPLEMENTARY TABLE 2**. In all cases following amplification of a plasmid or gene, the amplicons were cleaned up with a DNA Clean & Concentrator - 5 kit from Zymo Research (cat. D4013) before cloning. Following cloning in all situations, ∼3µL of each cloning reaction was then transformed into 5-alpha Competent *Escherichia coli* cells from NEB (cat. C2987H), plated onto LB media containing AMP (100µg/mL) and NAT (250µg/mL), and left to grow at 30°C static for a day. Individual transformants were then patched onto LB + AMP + NAT agar plates and grown at 30°C static for an additional day. The original vector used in this study was our *C. albicans* CRISPRi-dCas9 plasmid (pRS159, Addgene #122378) [56].

To construct our dCas12 plasmid, we first used the IDT Codon Optimization Tool to generate the sequence for a *C. albicans* codon-optimized nuclease-deficient (d)Cas12 from *Lachnospiraceae* bacterium ND2006 (LbCas12a) previously employed in *Saccharomyces cerevisiae* [59]. The *C. albicans* dCas12a and the VPR construct from our *C. albicans* CRISPRa-dCas9 plasmid (pRS156, Addgene #182707) [57] were synthesized and inserted sequentially into pRS159 in place of dCas9-Mxi1 by Twist Bioscience, resulting in pRS835. The artifact dCas9 sgRNA scaffold was then removed via Gibson assembly with gRS89 (a donor repair template lacking the dCas9 sgRNA scaffold) and the NEBuilder® HiFi DNA Assembly Master Mix (cat. E2621L), resulting in pRS1003. To generate the four amino acid mutations for hyperdCas12a (D156R, D235R, E292R, D350R) [54], we synthesized a *C. albicans* codon-optomized gBlock from IDT (gRS104) encompassing the relevant region of DNA with the four amino acid replacements, and cloned it in place of the existing region via inverse PCR with the repliQa HiFi ToughMix (cat. 95200-025) and Gibson assembly as before, resulting in pRS1005, the final constitutively-expressed hyperdCas12a-CRISPRa plasmid backbone.

To generate the CRISPRi-dCas12a plasmid, we amplified the entirety of pRS1005 without VPR, and then amplified the Mxi1 repressor from pRS159, both with the repliQa HiFi ToughMix. By adding overhanging regions of homology in the primers we used to amplify Mxi1, we were then able to clone the Mxi1 construct into the amplified pRS1005 via Gibson assembly, resulting in pRS1074, the constitutively-expressed hyperdCas12a-CRISPRi plasmid. Finally, to replace the *ACT1* promoter that drives the expression of dCas12a in both pRS1005 and pRS1074 with the Tet-Off system [71], we amplified the entirety of both plasmids omitting *pACT1*, and amplified the Tet-Off promoter construct from our inducible CRISPRi-dCas9 system (pRS479, Manuscript in Preparation). We then cloned the linearized vectors together with the Tet-Off construct again via Gibson assembly as described above, resulting in pRS1073 (CRISPRa) and pRS1211 (CRISPRi). As the TetO construct contains the recognition site of PmlI (thereby making the PmlI site near the gRNA construct no longer unique), we then amplified the entirety of pRS1073 and pRS1211 around a region upstream of the *SNR52* promoter with the repliQa HiFi ToughMix, and cloned in an additional NaeI site using a single-stranded oligo insert via the NEBuilder® HiFi DNA Assembly Master Mix once again. This resulted in pRS1242 (CRISPRa inducible) and pRS1244 (CRISPRi inducible).

Whole Plasmid Sequencing was performed by Plasmidsaurus using Oxford Nanopore Technology with custom analysis and annotation for clones at each step of the plasmid construction process. All five final plasmids have been deposited to AddGene: CRISPRa-dCas12a constitutive (pRS1003, AddGene ID: 247640), CRISPRa-hyperdCas12a constitutive (pRS1005, AddGene ID: 247641), CRISPRi-hyperdCas12a constitutive (pRS1074, AddGene ID: 247642), CRISPRa-hyperdCas12a inducible (pRS1242, AddGene: 247643), and CRISPRi-hyperdCas12a inducible (pRS1244, AddGene: 247644).

### crRNA design, synthesis, and cloning

crRNAs were designed using the Eukaryotic Pathogen CRISPR guide RNA/DNA Design Tool with the *C. albicans* SC5314 FungiDB-26 genome selected, and with a custom PAM of “TTTV” on the 5’ end [90]. Generally, a region around 90-370bp upstream of the start codon of the target gene was considered when designing crRNAs. To generate the non-targeting crRNAs and confirm they had no matches in the *C. albicans* genome, random 20bp sequences were first created and BLASTed against the *C. albicans* SC5314 Assembly 22 genome using the *Candida Genome Database* BLAST tool [89]. The direct repeat sequence used for singeplexed crRNA arrays was “taatttctactaagtgtagat”, and the direct repeat sequence used for multiplexed crRNA arrays was “aatttctactaagtgtagat” (**SUPPLEMENTARY TABLE 3** and **SUPPLEMENTARY TABLE 4**), similarly to what has been reported previously [59].

For single-crRNA arrays, each array was ordered as a forward and reverse-complement single-stranded DNA oligo pair from Integrated DNA Technologies (IDT). Oligos were resuspended in Nuclease Free Duplex Buffer from IDT (cat. 11-05-01-12) to 100µM, heated at 94°C for one minute, mixed with their counterpart oligo in equal volumes, and heated again at 94°C for two minutes. The duplexed oligo was then cloned into the corresponding plasmid via a Golden Gate strategy with the following mix: ∼1000ng of plasmid, 1µL of duplexed oligo, 2µL of 10X rCutSmart buffer, 2µL of ATP, 1µL of SapI, 1µL of T4 DNA ligase, and nuclease-free water to a final volume of 20µL. Reactions were incubated in a thermocycler using the following conditions: (37°C for 2 min, and 16°C for 5 min) for 99 cycles, 65°C for 15 min, and 80°C for 15 min. Following this, an additional 1µL of SapI was added to each reaction and incubated at 37°C for 1h to remove any plasmids not containing the properly cloned crRNA.

For multiple-crRNA arrays, insert arrays were designed with ∼20bp of homology to the receiving vector on either end, and ordered as GenTitan Gene Fragments from GenScript. They were then resuspended in 30µL of nuclease-free water and quantified via NanoDrop using a Infinite 200 PRO microplate reader (Tecan). Plasmids were either treated with both NaeI (cat. R0190S) and PmlI (cat. R0532S) for the constituvely-expressing plasmids, or NaeI alone for the inducible plasmids, in all cases involving ∼1000ng of vector, 1µL of corresponding enzyme, 5µL of 10X rCutSmart Buffer, and nuclease-free water to a total volume of 50µL, where reactions were incubated at 37°C for 12h followed by a 20 min 65°C heat-inactivation step. Digested plasmids were then cleaned up with a DNA Clean & Concentrator - 5 kit from Zymo Research (cat. D4013). Next, 100ng of cleaned-up vector was mixed with a corresponding gene fragment containing the insert CRISPR array at a 3:1 ratio, along with 10µL of NEBuilder® HiFi DNA Assembly Master Mix as before, and nuclease-free water to a total volume of 20µL. Control reactions set up in the same way but with nuclease-free water instead of an insert were also established in each case to allow for the proportion of false-positive transformants to be determined. Finally, reactions were incubated at 50°C for 30 min.

For cells transformed with a plasmid containing only one crRNA, transformant patches were PCR tested for the presence of the corresponding crRNA as previously described [91], where one primer in the PCR is one of the single-stranded crRNA oligos, therefore allowing for a direct test of the presence of the array. For cells transformed with a plasmid containing a multiple-crRNA array, a region encompassing the entire crRNA array was amplified and sent for Sanger sequencing to verify that the gene fragment was correctly integrated into the plasmid. Overnight cultures of verified transformant patches were made in 5mL of LB with AMP and NAT and grown at 30°C (∼250 RPM), and plasmids were then miniprepped with a GeneJET Plasmid Miniprep Kit from ThermoFisher (cat. K0503) as per the manufacturer’s instructions.

### Candida albicans transformation

Fungal transformations were performed as previously described with some minor modifications [91]. *C. albicans* cells were grown up overnight in YPD at 30°C (∼250 RPM). Plasmids were linearized using PacI from NEB (cat R0547L), and left at 37°C for 12h followed by a 20 min heat-inactivation step at 65°C. A transformation master mix was prepared with 800µL of 50% polyethylene glycol (PEG), 100µL of 10X Tris-EDTA buffer solution, 100µL of 1M lithium acetate, 40µL of UltraPure™ Salmon Sperm DNA Solution from ThermoFisher (cat. 15632-011), and 20µL of 1M dithiothreitol (DTT). Pelleted C. albicans cells were then resuspended in the transformation mix and the linearized plasmid, and then the reactions were left to incubate at 30°C for 1h. The solutions were then heat shocked at 42°C for 50 min in a water bath. Pelleted heat-shocked cells were washed with 1mL of fresh YPD three times, then transferred into tubes with 9mL of YPD (for a total of 10mL) and incubated for 4h at 30°C (∼250 RPM) to allow for expression of the NATr construct. Transformed cells were then pelleted, resuspended in ∼200µL of YPD, and plated on YPD + NAT (250µg/mL). Plates were grown at 30°C for 2 days, and a few transformants were then patched onto plates of YPD + NAT (250µg/mL) and grown at 30°C for an additional day. For cells transformed with a plasmid containing only one crRNA, the patches were PCR tested for the presence of the corresponding crRNA again, as described above. For cells transformed with plasmids containing a multiple-crRNA array, the entire crRNA array was amplified, and transformants were selected by size for the correct plasmid.

### Growth curve assays

For all growth-curve assays, the well position of each strain and condition were randomized to avoid any positional bias for growth in different areas of the plate. When possible, the wells of the outer ring of the plate were only filled with 200μL of blank YPD to avoid potential edge effects. For experiments comparing the growth of all NTCs and the WT, overnight fungal cultures grown in YPD at 30°C were each diluted to an OD600 of 0.05 in 1mL of fresh YPD. The diluted cultures were then added in 200μL aliquots to four randomized wells in a 96-well flat-bottomed plate. To the remaining empty wells in the 96-well plate, 200μL of blank YPD was added. For experiments comparing the growth of CRISPRi strains targeting *ERG6* and *ERG251*, stocks of fluconazole were prepared to twice as high as the desired concentration, and aliquoted out in 100μL volumes into the corresponding wells. Overnight cultures were first diluted to an OD600 of 0.1 in 1mL of YPD, and 100μL of each was then further diluted by 60-fold (for example, 50μL into 3mL of YPD). 100μL of each final diluted culture was then mixed into the corresponding wells in quadruplicate with the varying concentrations of fluconazole that were previously added, for a final OD600 of ∼0.000833, and a final volume in each well of 200μL. In all cases, 30μL of mineral oil (cat. MIN333.500, BioShop) was then added to the top of each well. Growth was measured at 37°C via optical density at 600nm every 15 minutes for 24-48h using an Infinite 200 PRO microplate reader (Tecan), where plates were shaken orbitally for 800s at a 4mm amplitude in between growth measurements. Area-under-the-curve (AUC) values were calculated by using the trapezoidal rule using a custom Python script. Raw growth curve data are listed in **SUPPLEMENTARY DATA 1**.

### Minimum inhibitory concentration assays, and IC_50_ and SMG calculations

MIC assays were performed in 96-well flat-bottomed plates, where 200μL of drug stock diluted in YPD to twice the highest concentration desired was first added to each well in the first column. The drug was then serially diluted ½ in YPD over the subsequent columns until the second-to-final column. Overnight cultures of *C. albicans* grown in YPD at 30°C (∼250 RPM) were then diluted to an OD600 of 0.1 in 1mL of YPD, and 100μL of each was then further diluted by 60-fold (for example, 50μL into 3mL of YPD). 100μL of each final diluted culture was then mixed into one row each of the 96-well plate, for a final OD600 of ∼0.000833. Each plate contained one row of WT SC5314 *C. albicans* as a control, and a row of blank media as a contamination control. All strains were tested in triplicate, and plates were read following incubation at 37°C (static) for 24h and 48h, unless otherwise stated. The MIC assays in Figure 2F were performed in parallel on the same day, but split up into two sets of 96-well plates. Therefore, the NTC values shown in the corresponding wells in Figure 2F represent an average between the two sets of triplicate plates. OD600 values were read using an Infinite 200 PRO microplate reader (Tecan). IC_50_ values were calculated by fitting a logistic dose-response curve to the normalized OD values via a custom Python script. SMG values were calculated as has been previously described with some minor modifications [92]. First, we identified the MIC_80_ for each strain replicate by dividing each well to the corresponding no-drug well, and checking to see if the resulting value was above 80 after multiplying by 100. We opted to use the MIC_80_ instead of the MIC_50_ since some of the highly tolerant strains would have otherwise been called resistant despite clearly showing a drop in the OD600 at a certain concentration of drug (**SUPPLEMENTARY DATA 2**). Following this, we averaged the OD600 values of all wells that were above the MIC_80_ and divided them by the corresponding no-drug well to produce the SMG value. The statistical significance of the IC_50_ and SMG values were calculated using an unpaired, two-tailed, parametric Welch’s t-test with GraphPad Prism version 10.5.0 for macOS, GraphPad Software, Boston, Massachusetts USA, www.graphpad.com. Raw MIC data are listed in **SUPPLEMENTARY DATA 2**.

### RNA extraction and reverse transcription quantitative PCR

Overnight cultures of strains for RT-qPCR were first diluted in triplicate to an OD600 of 0.05 in 40mL of YPD and left to grow at 240RPM for ∼4 hours, such that the OD600 values were above 0.2. Subculturing was performed at either 30°C or 37°C, depending on the experiment, as specified in the Results section. For the experiments verifying the inducibility of the systems only, overnight cultures were first grown in the presence of 250ng/mL anhydrotetracycline hydrochloride (ATc) from Alfa Aesar (cat. J66688). The following day, strains were then diluted in triplicate to an OD600 of 0.05 in 40mL of YPD with either 250ng/mL ATc or the corresponding volume of DMSO, and left to grow at 240RPM for ∼5 hours, such that the OD600 values were above 0.2. In all cases, following this growth period, the cultures were pelleted, re-suspended in ∼1mL of YPD, and frozen at -80°C. RNA extractions were performed using the PuroSPIN™ Total RNA Purification Plus Kit from Luna Nanotech (cat. NK251-200) with an enzymatic digestion of ∼300μL of thawed pelleted cultures. Briefly, pelleted cultures were resuspended in a solution of 1M sorbitol and 0.1M EDTA adjusted to pH 7.4, 0.1% β-mercaptoethanol, and ∼200 units of Zymolase from BioShop (cat. ZYM001.1). The cell solutions were then incubated at 30°C for around 45 minutes with gentle agitation before being centrifuged at 4000 X G for 5 minutes to pellet the spheroplasts. After the supernatants were discarded, 400μL of the Working Buffer LB-R (supplemented with 20μL of 14 M β-mercaptoethanol per 1mL) was added, and the spheroplasts were vortexed thoroughly for 1-2 minutes. All steps following this were done exactly as per the manufacturer’s instructions. RNA was quantified via NanoDrop using an Infinite 200 PRO microplate reader (Tecan). Preliminary testing of RNA extracted with the PuroSPIN™ Total RNA Purification Plus Kit in this way on TapeStation revealed RIN values approaching 10.0. RNA was stored at -80°C if not used immediately.

RT-qPCR was performed with the Luna® Universal One-Step RT-qPCR Kit from NEB (cat. E3005S) using an input of 200-300ng of RNA (exact amount was kept consistent within each experiment). RNA was first diluted to 40ng/μL, and 5μL was mixed into a corresponding well on a qPCR 96 well Hard Shell Plate from Applied Biosystems (cat. 4483354) with 15μL of the master mix prepared as per the Luna® Universal One-Step RT-qPCR Kit with the corresponding RT-qPCR primers, which are listed in **SUPPLEMENTARY TABLE 5**. Reactions were performed primarily using a QuantStudio™ 3 Real-Time PCR system from Applied Biosystems. Fold-changes in gene expression were calculated via the comparative C_T_ method [93]. Briefly, the Cq value of the gene of interest in each RNA sample was subtracted from the housekeeping gene *PMA1* (unless otherwise specified) to obtain a _Δ_Ct value [94]. The _Δ_Ct values in the experimental CRISPR-dCas strains were then compared to the corresponding non-targeting control strain (or control condition in the case of the inducible systems) to obtain a _ΔΔ_CT value and finally a fold difference in expression of the target gene(s). The statistical significance when comparing differences in fold-change in target gene expression between strains was calculated using an unpaired, two-tailed, parametric Welch’s t-test with GraphPad Prism version 10.5.0 for macOS, GraphPad Software, Boston, Massachusetts USA, www.graphpad.com. Raw RT-qPCR data are listed in **SUPPLEMENTARY DATA 3**.

## Acknowledgements

This work was supported by a a Natural Sciences and Engineering Research Council of Canada (NSERC) Discovery Grant (RGPIN-2018-4914). NCG is supported by an NSERC CGS-D award, and RSS is supported by a Canada Research Chair in Microbial Functional Genomics and Synthetic Biology.

## Author Contributions

NCG, RKJR, and MRR performed the experiments. NCG designed the experiments, analyzed the data, generated the figures, and drafted the manuscript. NCG and RSS conceptualized the project and revised the manuscript together. RSS supervised the project, provided resources, and acquired funding.

## Supplementary Figures

**Supplementary Figure 1.**
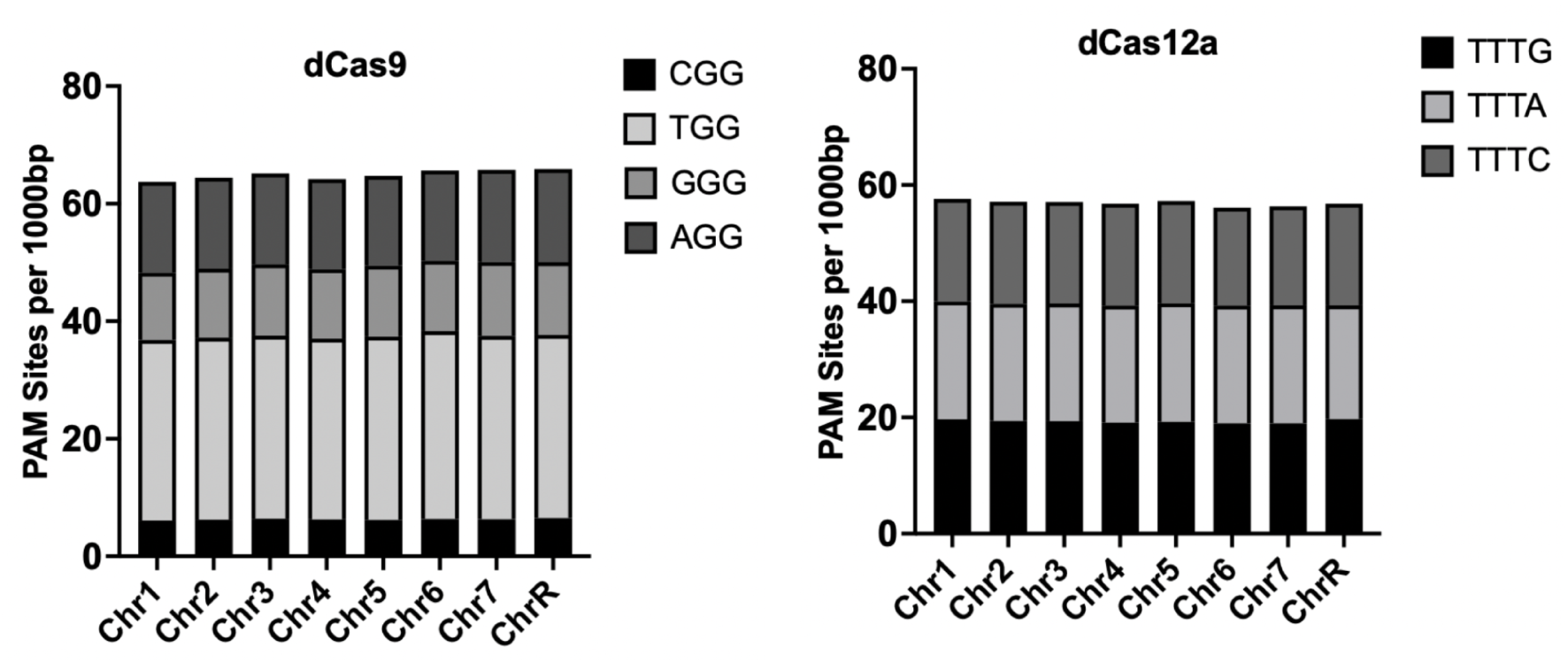
Canonical PAM sites available for dCas9 and dCas12a in each chromosome of *C. albicans,* normalized by size. PAM sites were searched across the *C. albicans* genome by using a custom Python script. The total number identified on each chromosome was then divided by the number of base pairs (bp) in the corresponding chromosome, and multiplied by 1000 to get PAM sites per 1000bp.

**Supplementary Figure 2.**
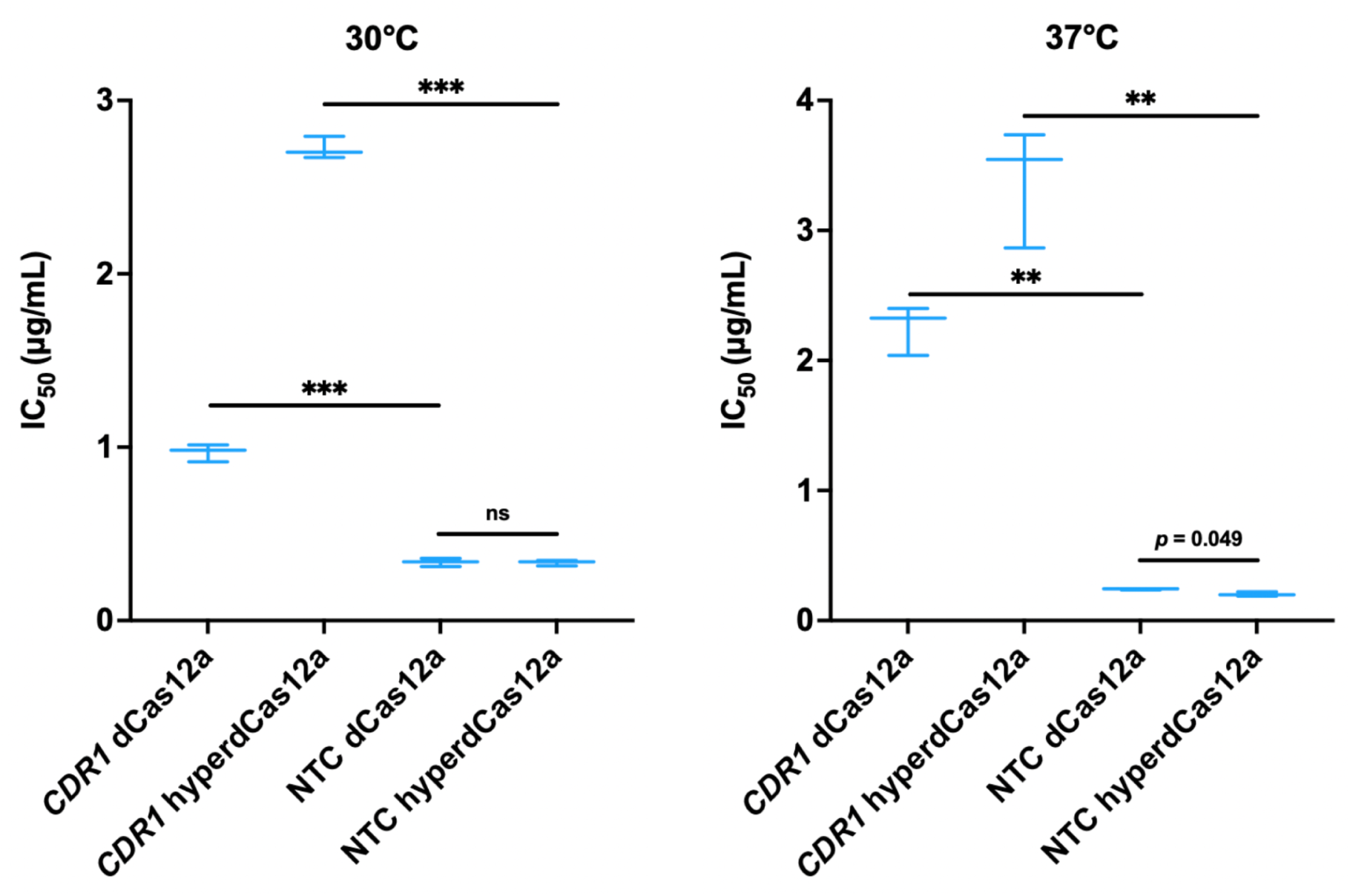
The different levels of overexpression of *CDR1* achieved via dCas12a and hyperdCas12a lead to an expected corresponding increase in fluconazole IC_50_ values. Strains were grown in an MIC assay at 37°C in YPD with a concentration gradient of fluconazole, and OD600 values were measured after 24 hours of growth. A logistic dose-response curve was then fitted to the normalized OD600 values from the MIC data to obtain the IC_50_ values. Statistics were performed with an unpaired, two-tailed, parametric Welch’s t-test, ns *= p* > 0.05, \*\**p* < 0.01, \*\*\**p* < 0.001.

**Supplementary Figure 3.**
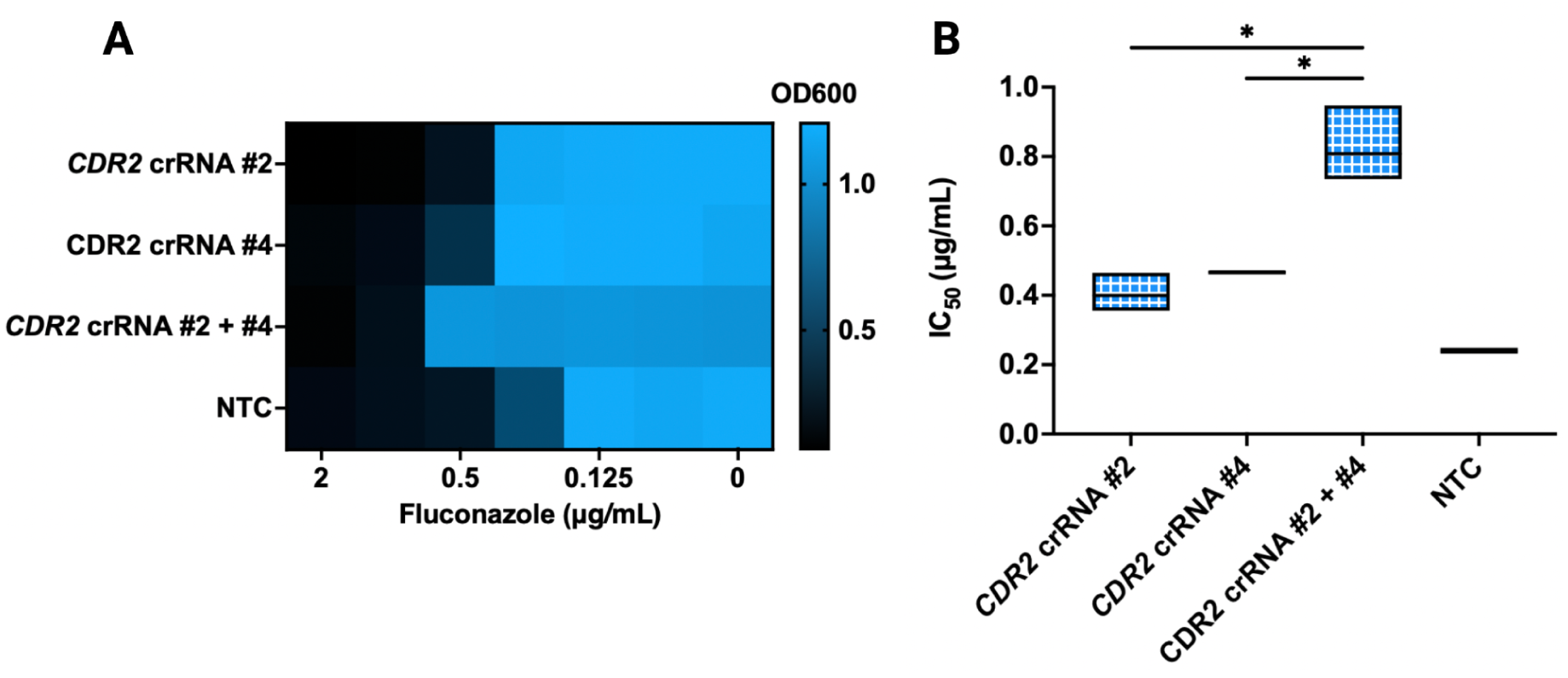
The enhanced overexpression of *CDR2* from tiling the promoter with two crRNAs results in increased fluconazole resistance. (**A**) Fluconazole MIC assay with the single-crRNA and multiple-crRNA *CDR2* CRISPRa strains. Strains were grown at 37°C in YPD with a concentration gradient of fluconazole, and OD600 values were measured after 24 hours of growth. (**B**) Fluconazole IC_50_ values of the single-crRNA and multiple-crRNA *CDR2* CRISPRa strains. A logistic dose-response curve was fitted to the normalized OD600 values from the MIC assay in SUPPLEMENTARY FIGURE 3A to obtain the IC_50_ values. Statistics were performed with an unpaired, two-tailed, parametric Welch’s t-test, \**p* < 0.05.

## Tables

**TABLE 1:**
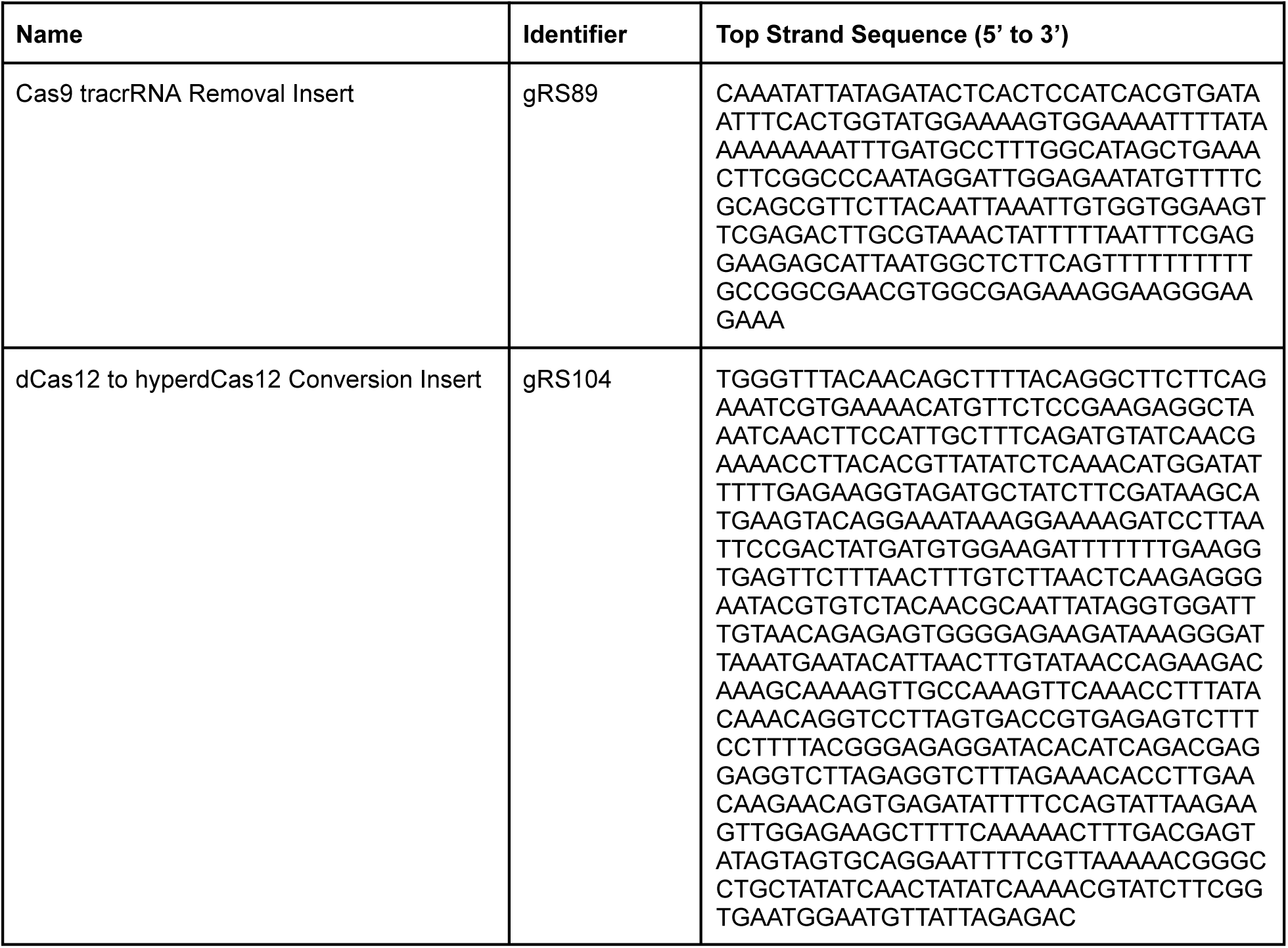
Double-stranded gene fragments for cloning used in this study.

**TABLE 2:**
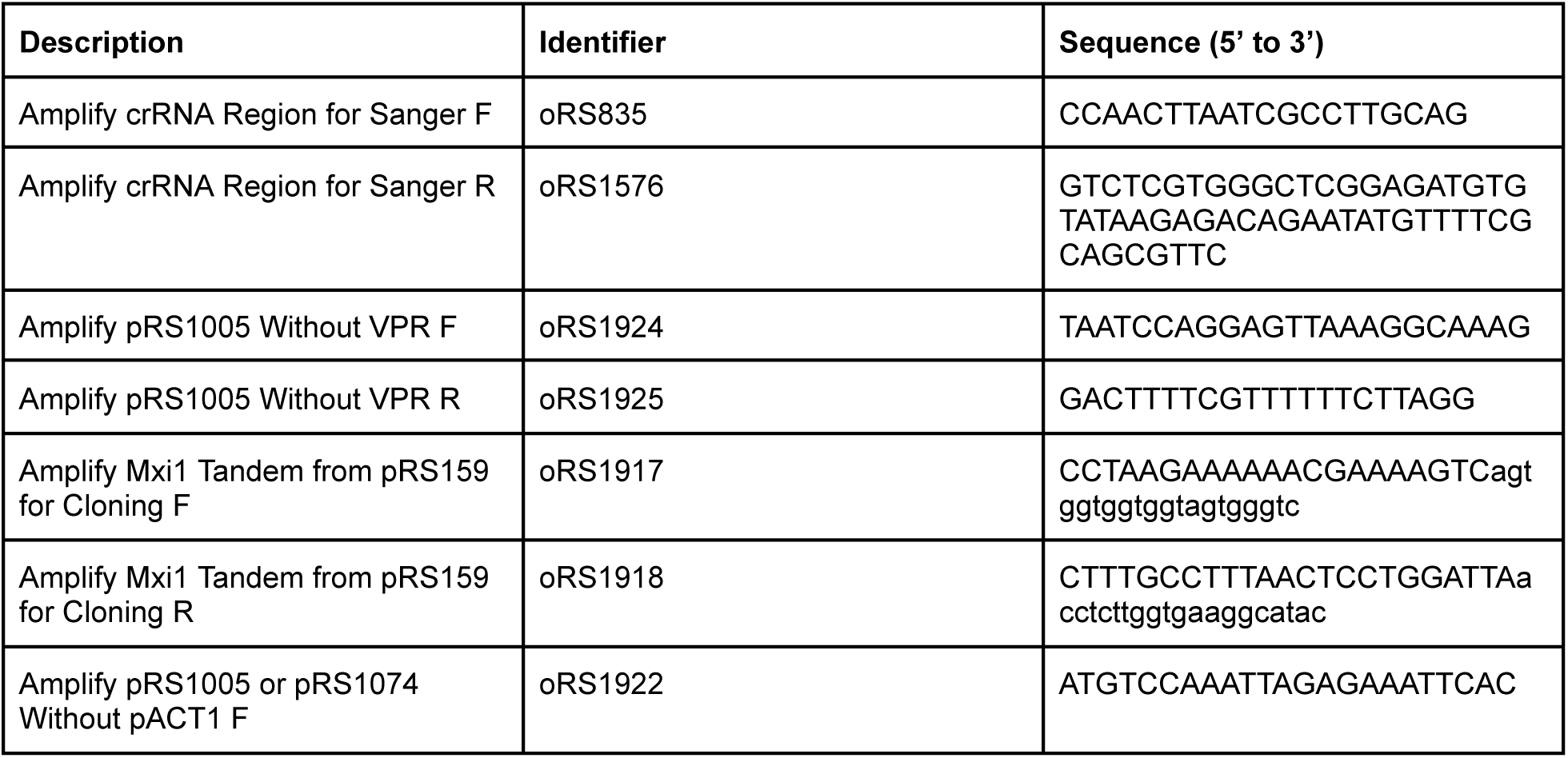
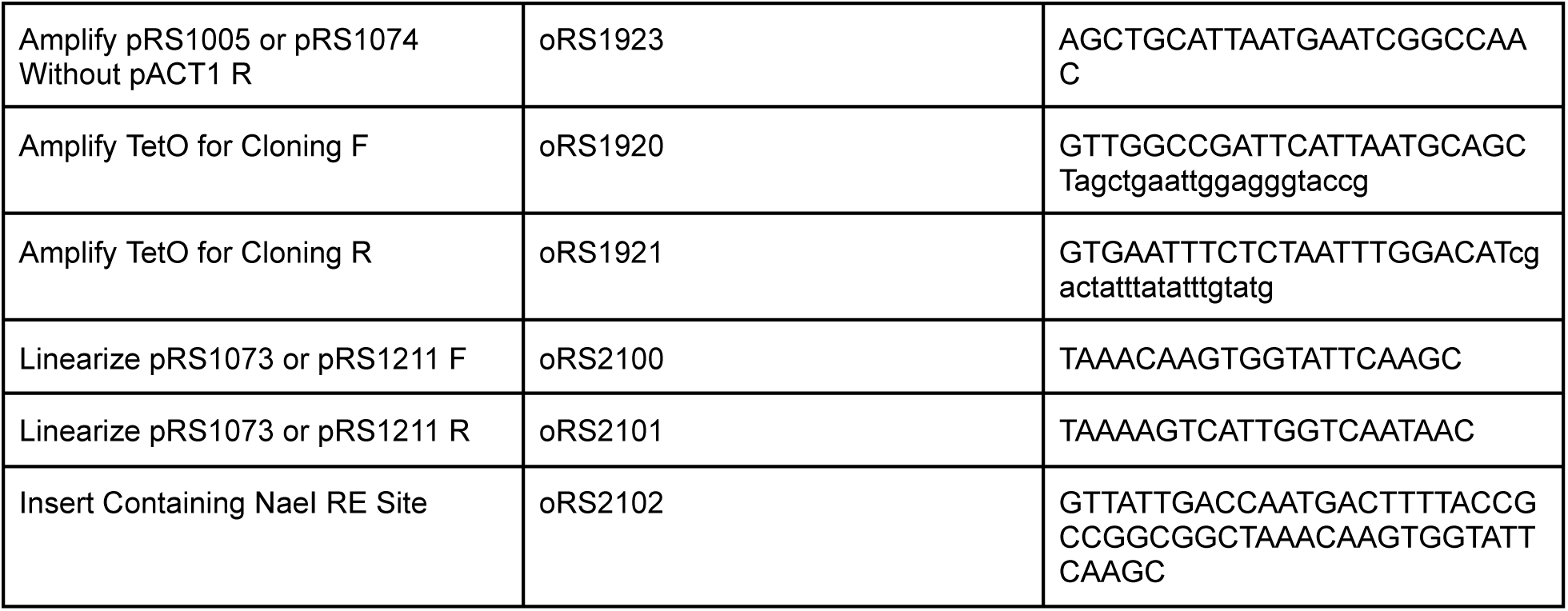
Primers/single-stranded gene fragments for cloning used in this study.

**TABLE 3:**
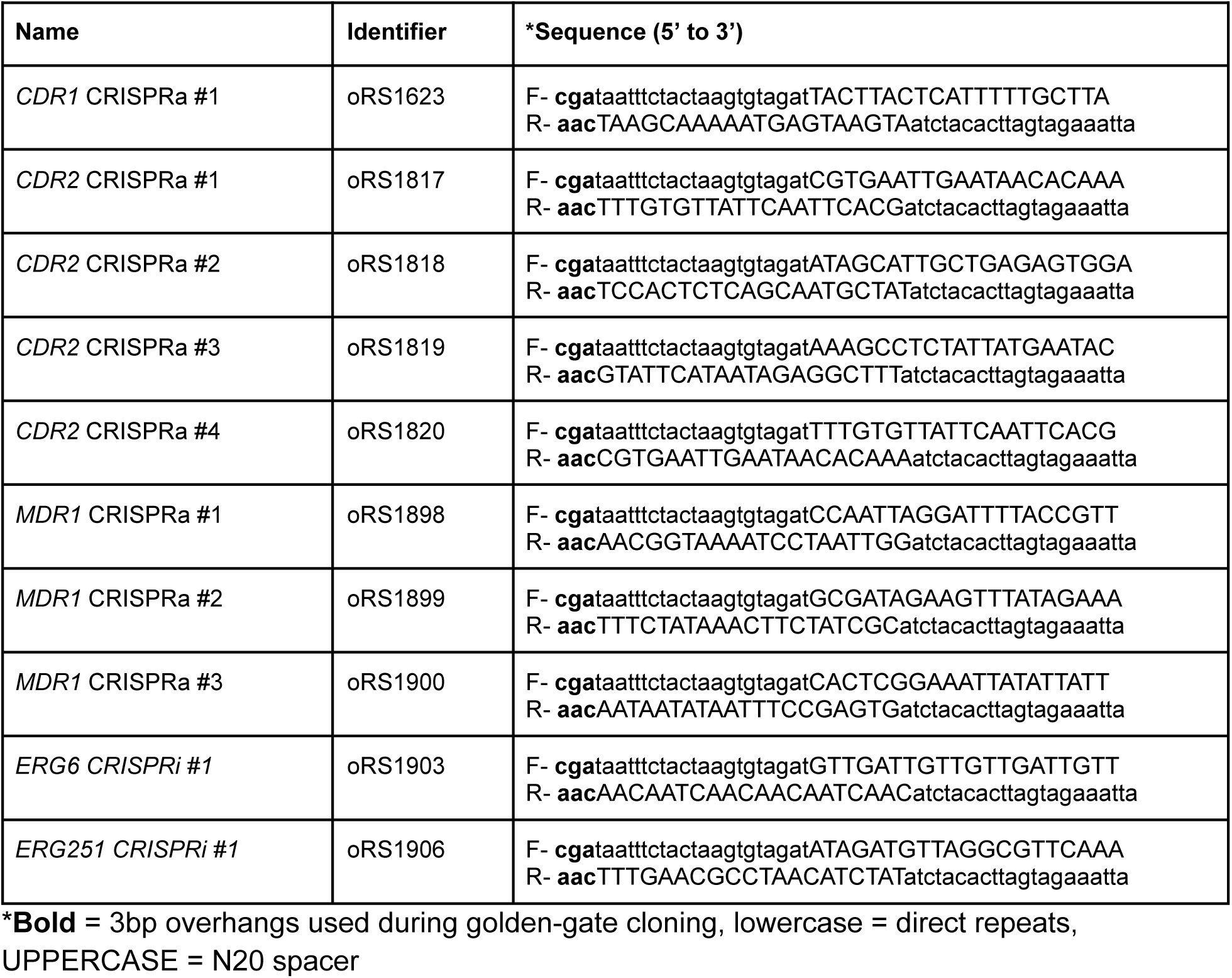
crRNA sequences for singleplexed targeting used in this study.

**TABLE 4:**
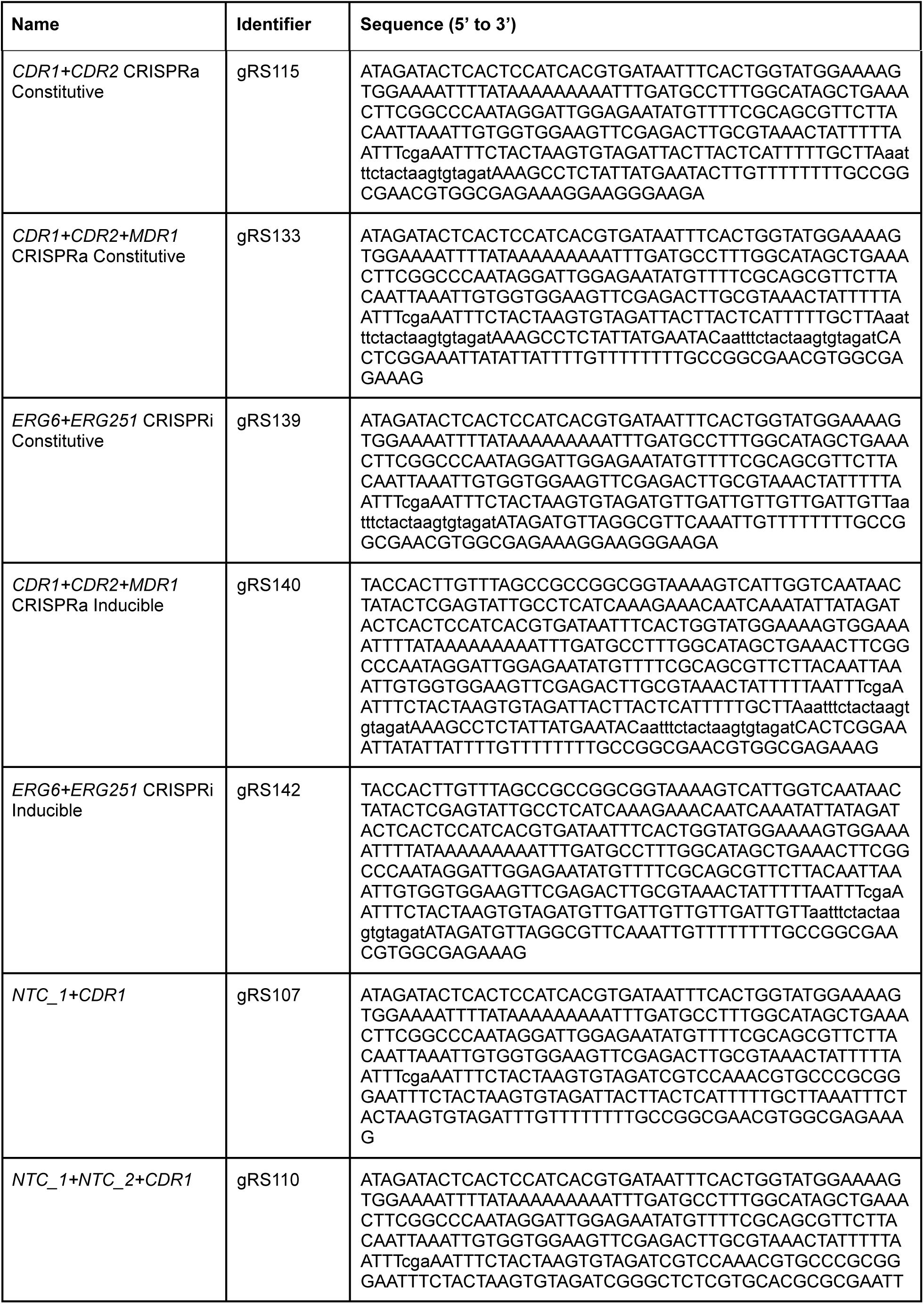
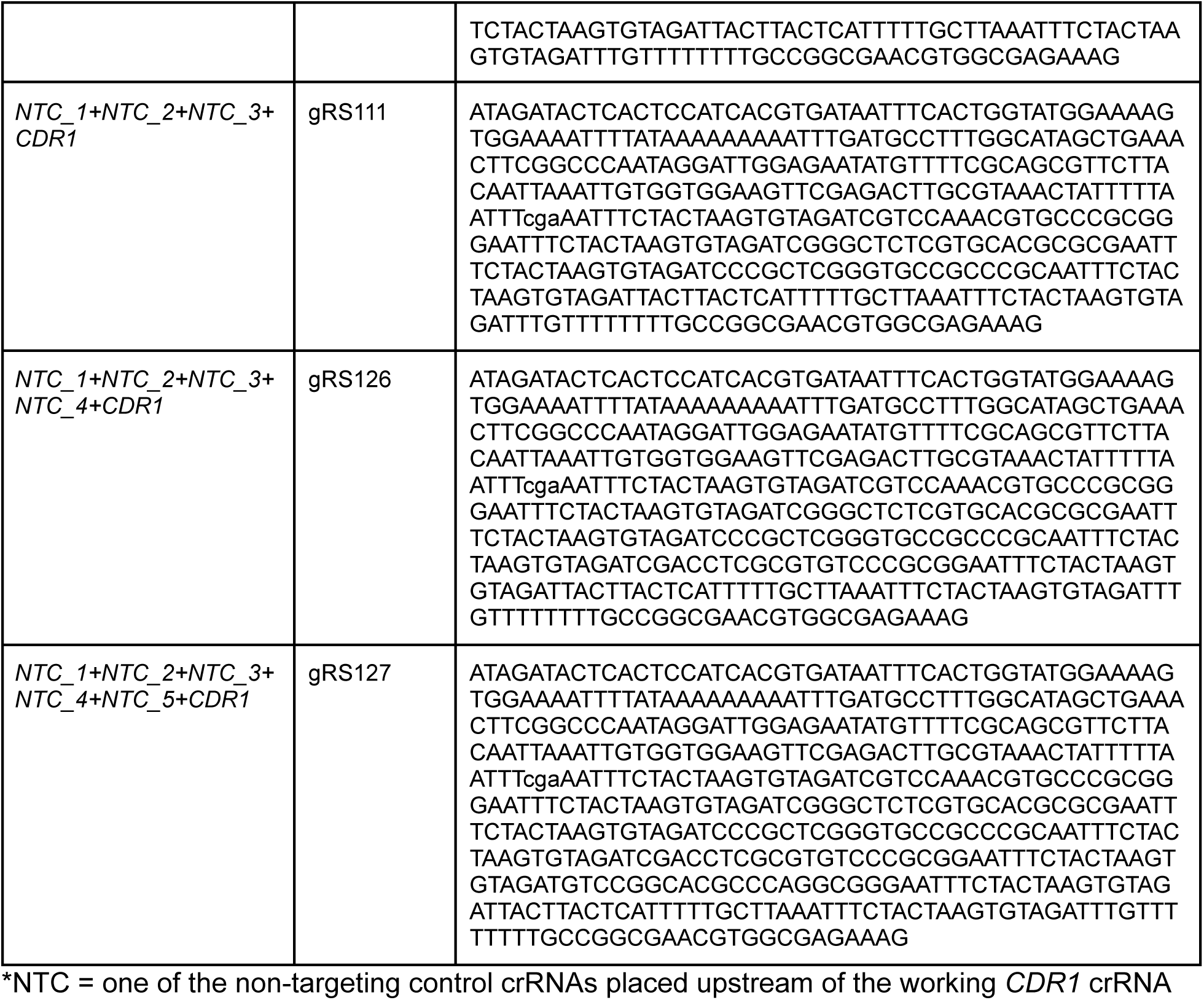
crRNA array DNA fragments for multiplexed targeting used in this study.

**TABLE 5:**
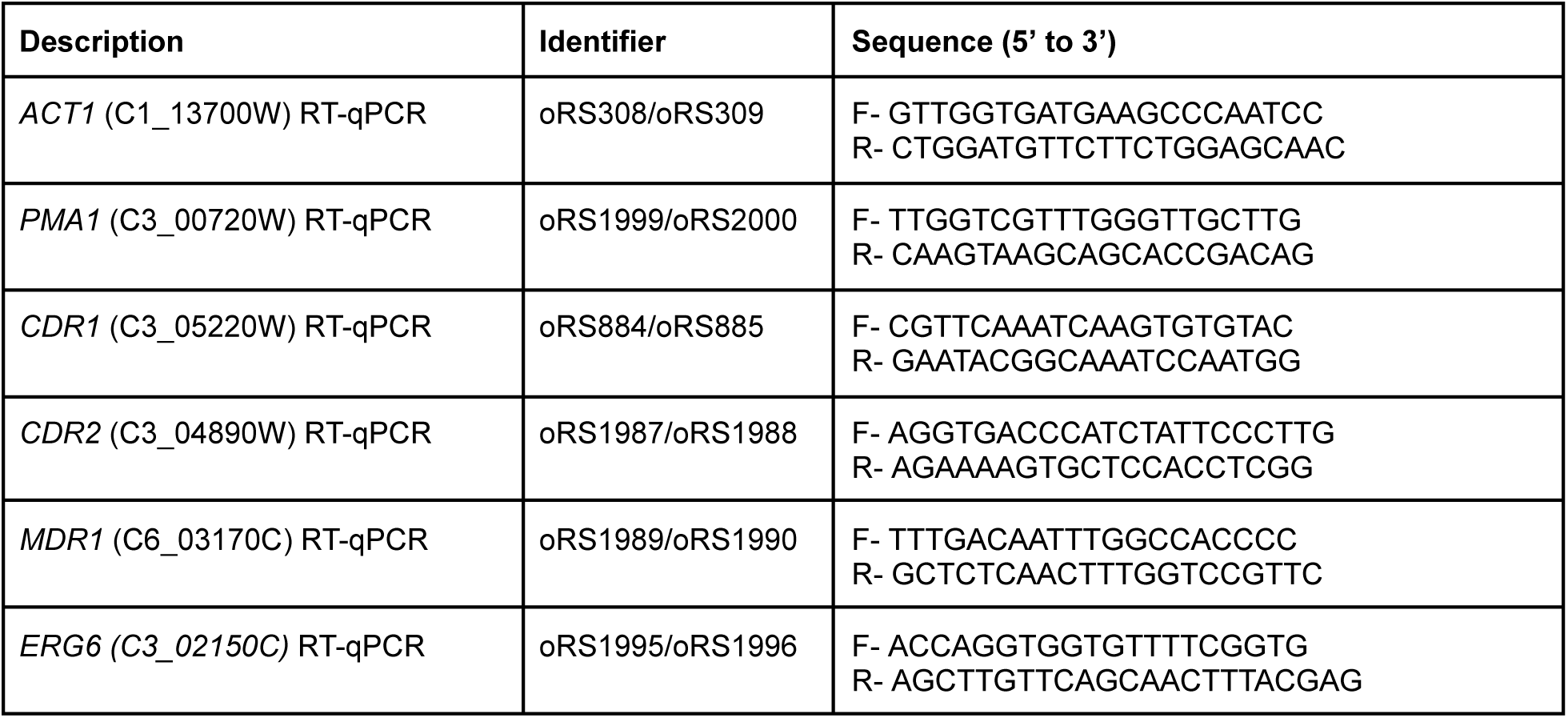
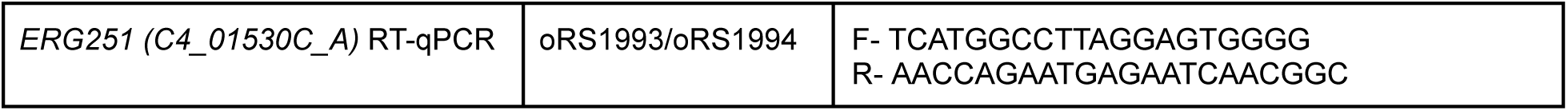
RT-qPCR primers used in this study.

**TABLE 6:** Plasmids used in this study.

| Description | Identifier | AddGene |
| --- | --- | --- |
| CRISPRa-dCas12a Constitutive with Artifact dCas9 sgRNA Scaffold | pRS835 | N/A |
| CRISPRa-dCas12a Constitutive | pRS1003 | TBD |
| CRISPRa-hyperdCas12a Constitutive | pRS1005 | TBD |
| CRISPRi-hyperdCas12a Constitutive | pRS1074 | TBD |
| CRISPRa-hyperdCas12a Inducible without Additional NaeI Site | pRS1073 | N/A |
| CRISPRi-hyperdCas12a Inducible without Additional NaeI Site | pRS1211 | N/A |
| CRISPRa-hyperdCas12a Inducible | pRS1242 | TBD |
| CRISPRi-hyperdCas12a Inducible | pRS1244 | TBD |

## Notes

### Competing Interest Statement

The authors have declared no competing interest.

